# Contrasting action and posture coding with hierarchical deep neural network models of proprioception

**DOI:** 10.1101/2020.05.06.081372

**Authors:** Kai J. Sandbrink, Pranav Mamidanna, Claudio Michaelis, Matthias Bethge, Mackenzie Weygandt Mathis, Alexander Mathis

## Abstract

Biological motor control is versatile and efficient. Muscles are flexible and undergo continuous changes, requiring distributed adaptive control mechanisms. How proprioception solves this problem in the brain is unknown. The canonical role of proprioception is representing the body state, yet we hypothesize that the proprioceptive system can decode high-level, multi-feature actions. To test this theory, we pursue a task-driven modeling approach.We generated a large synthetic dataset of human arm trajectories tracing the alphabet in 3D space and use a musculoskeletal model plus modeled muscle spindle inputs to extract muscle activity. We then contrast two tasks, one character trajectory-decoding and another action recognition task that allows training of hierarchical models to decode position, or classify the character identity from the spindle firing patterns. Artificial neural networks could robustly solve these tasks, and the networks’ units show tuning properties akin to neurons in the primate somatosensory cortex and the brainstem. Remarkably, only the action-recognition trained, and not the trajectory decoding trained, models possess directional selective units (which are also uniformly distributed), as in the primate brain. Taken together, our model is the first to link tuning properties in the proprioceptive system at multiple levels to the behavioral level. We find that action-recognition, rather than the canonical trajectory-decoding hypothesis, better explains what is known about the proprioceptive system.

## Introduction

Proprioception is a critical component of our ability to perform complex movements, localize our body in space, and adapt to environmental changes (1–3). Our movements are generated by a large number of muscles and are sensed via a diverse set of receptors, most importantly muscle spindles, which carry highly multiplexed information (2, 4). For instance, arm movements are sensed via distributed and individually ambiguous activity patterns of muscle spindles, which depend on relative joint configurations rather than the absolute hand position (5, 6). Interpreting this high dimensional input (around 50 muscles for a human arm) of distributed information at the relevant behavioral level poses a challenging decoding problem for the central nervous system (6, 7). Proprioceptive information from the receptors undergoes several processing steps before reaching somatosensory cortex (3, 8, 9) - from the spindles that synapse in Clarke’s nucleus, to the brainstem, thalamus (3, 10), and finally to somatosensory cortex (S1). In cortex, a number of tuning properties have been observed, such as responsiveness to varied combinations of joints and muscle lengths (11, 12), sensitivity to different loads and angles (13), and broad and uni-modal tuning for movement direction during arm movements (14). The proprioceptive information in S1 is then hypothesized to serve as the basis of a wide variety of tasks, via its connections to motor cortex and higher somatosensory processing regions (1–3, 15, 16).

One key role of proprioception is to sense the state of the body—i.e., posture. This information subserves many other functions, from balance to motor learning. Thus, to gain insights into the computations of the proprioceptive system, we quantitatively compare two different goals in a task-driven fashion: a trajectory-decoding task, and an action recognition task (Figure 1). The trajectory-decoding task represents the canonical view of proprioception (9, 17). Alternatively, the role of the proprioceptive system could be to infer actions (i.e., complex sequences of postures). Our hypothesis is motivated by the observation that action segmentation would be an efficient way to represent complex behavior, and it could directly drive the “action map” in motor cortex (18).

**Figure 1.**
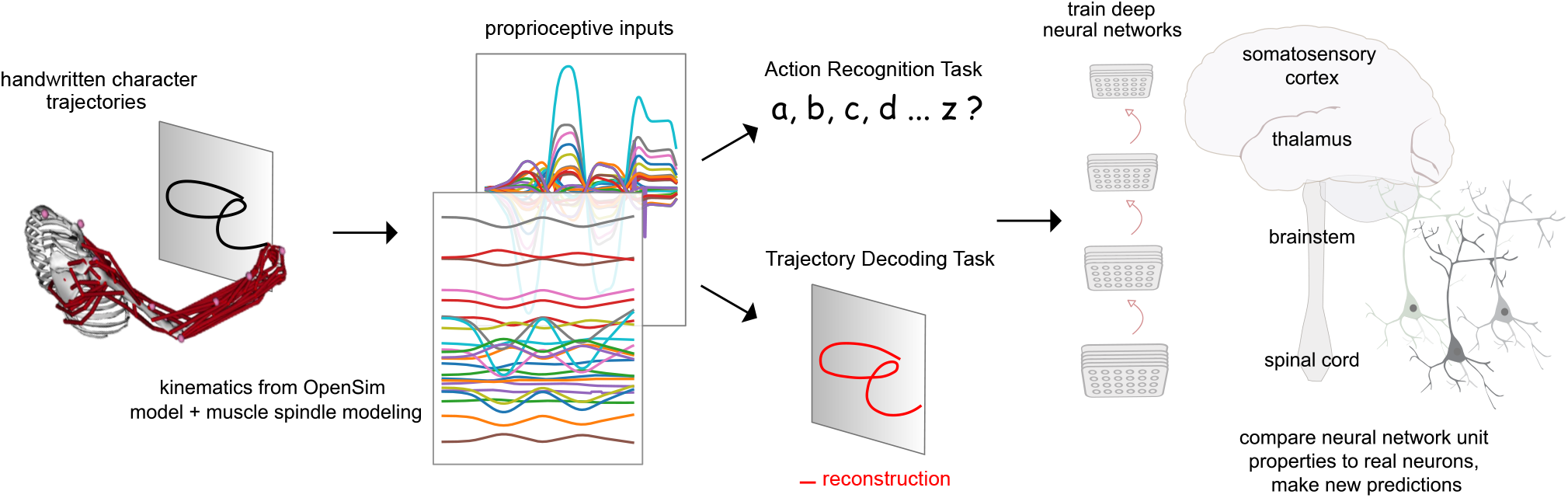
Contrasting spindle-based tasks to study proprioception: Proprioceptive inputs that correspond to the tracing of individual letters were simulated using a musculoskeletal model of a human arm. This scalable, large-scale dataset can then be used to train deep neural networks models of the proprioceptive pathway to either classify the character **(Action Recognition Task, ART)** or to localize the hand **(Trajectory Decoding Task, TDT)** based on the input muscle spindle firing rates. These models can be analyzed, compared and contrasted to what is known about the proprioceptive system in primates.

Large-scale datasets like ImageNet (19) that present a challenging visual object-recognition task, have allowed the training of deep neural networks whose representations closely resemble tuning properties of single neurons in the ventral pathway of primates (20–26). This goal-driven modelling approach (23, 27, 28) has since successfully been applied to other sensory modalities such as touch (29, 30), thermosensation (31) and audition (32). However, unlike for vision and audition, where large annotated datasets of raw images or sound are readily available, data of relevant proprioceptive stimuli (as well as task goals) are not.

To create a large-scale passive movement dataset, we started from human motion data for drawing different Latin characters (33). Next, a musculoskeletal model of the human upper limb (34) is employed to generate muscle length configurations corresponding to drawing the pen-tip trajectories in multiple horizontal and vertical planes. These are ultimately converted into proprioceptive inputs using models of spindle Ia and II models (35, 36). We then used the tasks to train families of neural networks to either decode the full trajectory of the hand-written characters, or classify the characters, both purely from the estimated spindle firing rates. Through an extensive hyper-parameter search we found neural networks for various architectures that optimally solve the task. We then analyzed those models and found that models trained on the action recognition, but not the trajectory decoding task more closely resemble what is known about tuning properties in the proprioceptive pathway. Collectively, we present a framework for studying the proprioceptive pathway using goal-driven modeling by synthesizing datasets of muscle (spindle) activities in order to test new theories of coding.

## Results

### Spindle-based biomechanical character recognition task

To model the proprioceptive system we designed two real-world proprioceptive tasks. The objective is to either classify or reconstruct Latin alphabet characters based on the proprioceptive inputs that arise when the arm is passively moved (Figure 1). In this way, we effectively computationally isolate proprioception from active movement—a challenge in experimental work. To create this we used a dataset of pen-tip trajectories for the 20 characters that can be handwritten in a single stroke (thus excluding *f, i, j, k, t* and *x*, which are multi-stroke) (33, 37). Then we generated one million end-effector (hand) trajectories by scaling, rotating, shearing, translating and varying the speed of each original trajectory (Figure 2A-C; Table 1).

**Figure 2.**
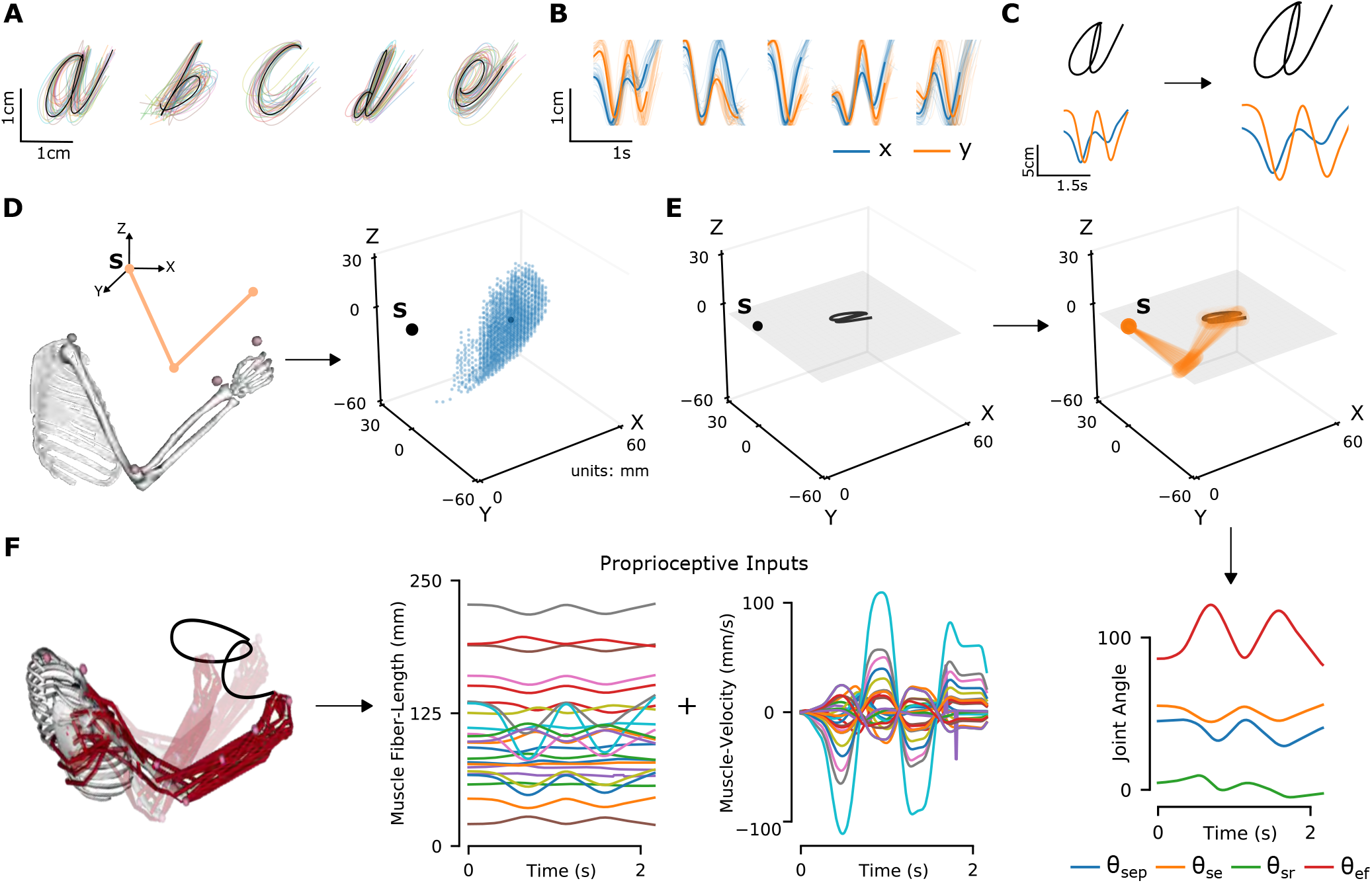
The proprioceptive character recognition dataset generation. (**A**) Multiple example pen-tip trajectories for five of the 20 letters are shown. (**B**) Same trajectories as in **A**, plotted as time courses of Cartesian coordinates. (**C**) Creating (hand) end-effector trajectories from pen-tip trajectories. (*left*) An example trajectory of character ‘a’ resized to fit in a 10cm x 10cm grid, linearly interpolated from the true trajectory while maintaining true velocity profile. (*right*) This trajectory is further transformed by scaling, rotating and varying its speed. (**D**) Candidate starting points to write the character in space. (*left*) A 2-link 4 degree of freedom (DoF) model human arm is used to randomly select several candidate starting points in the workspace of the arm (*right*), such that written characters are all strictly reachable by the arm. (**E**) (*left to right and down*) Given a sample trajectory in **C** and a starting point in the arm’s work-space, the trajectory is then drawn on either a vertical or horizontal plane that passes through the starting point. We then apply inverse kinematics to solve for the joint angles required to produce the traced trajectory. (**F**) (*left to right*): The joint angles obtained in **E** are used to drive a musculoskeletal model of the human arm in OpenSim, to obtain equilibrium muscle fiber-length trajectories of 25 relevant upper arm muscles. These muscle fiber-lengths and their instantaneous velocities together form the proprioceptive inputs.

**Table 1.**
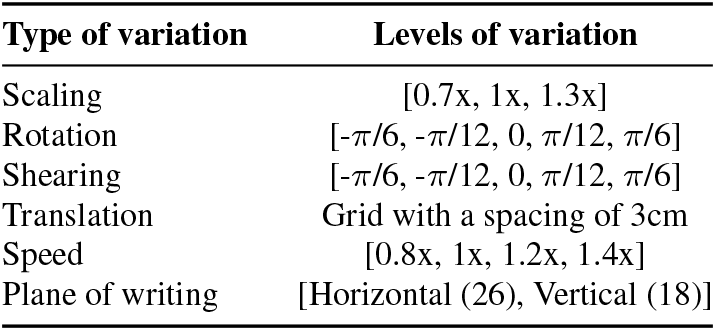
Variable range for the utilized data augmentation applied to the original pen-tip trajectory dataset. Furthermore, the character trajectories are translated to start at various starting points throughout the arm’s workspace. Overall yielding movements in 26 horizontal and 18 vertical planes.

To translate end-effector trajectories into 3D arm movements we computed the joint-angle trajectories through inverse kinematics using a constrained optimization approach (Figure 2D-E and methods). We iteratively constrain the solution space by choosing joint angles in vicinity of the previous configuration in order to eliminate redundancy. To cover a large 3D workspace we placed the characters in multiple horizontal (26) and vertical (18) planes and calculated corresponding joint-angle trajectories (starting points are illustrated in Figure 2D). A human upper-limb model in OpenSim (34) was then used to compute equilibrium muscle lengths for 25 muscles in the upper arm that lead to the corresponding joint angle trajectory (Figure 2F, Suppl. Video). We did not include hand muscles for simplicity, therefore the location of the end-effector is taken to be the hand location.

Based on these simulations, we generated proprioceptive inputs as muscle length and muscle velocity, which approximate receptor inputs during passive movement (see Methods). From this set, we selected a subset of two hundred thousand examples with smooth, non-jerky joint angle and muscle length changes, while making sure that the set is balanced in terms of the number of examples per class (see Methods). Since not all characters take the same amount of time to write, we padded the movements with static postures corresponding to the starting and ending postures of the movement and randomized the initiation of the movement in order to maintain ambiguity about when the writing begins. At the end of this process each sample consists of simulated proprioceptive inputs from each of the 25 muscles over a period of 4.8 seconds, simulated at 66.7 Hz. The dataset was split into a training, validation, and test set with a 72-8-20 ratio.

### Recognizing characters from muscle activity is challenging

We reasoned that several factors complicate the recognition of a specific character. Firstly, the end-effector position is only present as a distributed pattern of muscle activity. Secondly, the same character will give rise to widely different proprioceptive inputs depending on different arm configurations.

To test these hypotheses, we first visualized the data at the level of proprioceptive inputs by using t-distributed stochastic neighbor embedding (t-SNE, 38). This illustrated that character identity was indeed entangled (Figure 3A). Then, we trained pairwise support vector machine (SVM) classifiers as baseline models for character recognition. Here, the influence of the specific geometry of each character is notable. On average the pairwise accuracy is 86.6 ± 12.5 (mean ± S.D., *N* = 190 pairs, Figure 3B). As expected, similar looking characters were harder to distinguish at the level of the proprioceptive input—i.e. “*e*” and “*y*” were easily distinguishable but “*m*” and “*w*” were not (Figure 3B).

**Figure 3.**
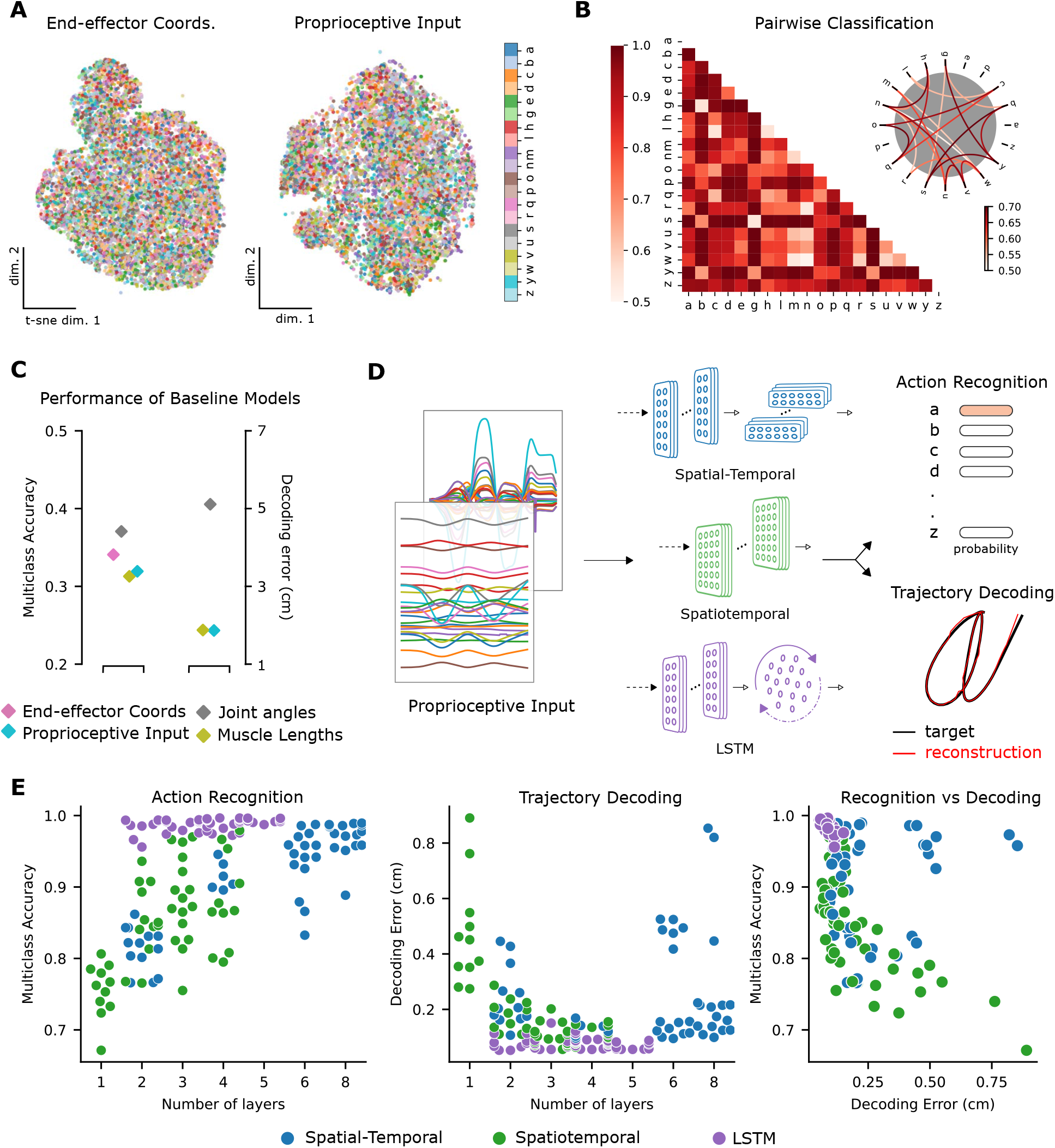
Quantifying action recognition and decoding task performance. (**A**) t-SNE embedding of the end-effector coordinates (left) and proprioceptive inputs (right). (**B**) Classification performance for all pairs of characters with binary SVMs trained on proprioceptive inputs. Chance level accuracy=50%. The pairwise accuracy is 86.6 *±* 12.5% (mean *±* S.D., *N* = 190 pairs). Subset of data is also illustrated as circular graph, edge color denotes the classification accuracy. For clarity, only pairs with performance less than 70% are shown, which corresponds to the bottom 12% of all pairs. (**C**) Performance of baseline models: Multi-class SVM performance computed using a one-vs-one strategy for different types of input/kinematic representations on the action recognition task (left). Performance of ordinary least-squares linear regression on the trajectory decoding task (right). Note there is no point for end-effector coordinates as this is trivial. (**D**) Neural networks are trained on two tasks - proprioceptive action recognition and trajectory decoding based on proprioceptive inputs. We tested three neural network architecture families. Each model is comprised of one or more processing layers as shown here. Processing of spatial and temporal information takes place through a series of 1-D or 2-D convolutional layers or a recurrent LSTM layer. (**E**) Performance of neural network models on the tasks: test performance of the 50 networks of each type is plotted against the number of layers of processing in the networks for the action recognition (left) and trajectory decoding (center) tasks separately, and against each other (right). Note we jittered the number of layers for visibility, but per model it is discrete.

To quantify the separability between all characters we used a one-against-one strategy with the trained pairwise classifiers (39). The performance of this multi-class decoder was poor regardless of whether the input was end-effector coordinates, joint angles, normalized muscle lengths, or proprioceptive inputs (Figure 3C). Taken together, these analyses highlight that it is difficult to extract the character class from those representations. Collectively, we demonstrated that the action recognition task is challenging as illustrated by t-SNE embedding (Figure 3A) and quantified by SVMs (Figure 3B, C). In contrast, as expected, accurately decoding the end-effector position (by linear regression) from the proprioceptive input is much simpler, with an average decoding error of 1.72cm, in a 3D workspace approximately 90×90×120cm^3^ (Figure 3C).

### Neural networks models of proprioception

We explore the ability of three types of artificial neural network models (ANNs) to solve the proprioceptive character recognition and decoding tasks. ANNs are powerful models for both their performance and for elucidating neural representations and computations (23, 40). An ANN consists of layers of simplified units (“neurons”) whose connectivity patterns mimic the hierarchical, integrative properties of biological neurons and anatomical pathways (23, 27, 41). As candidate models we parameterized a (spatio)temporal convolutional neural network, a spatial-temporal convolutional network (both TCNs, 42), and a recurrent neural network (a long short-term memory (LSTM) model, 43), which impose different inductive priors on the computations. We refer to these three types as spatial-temporal, spatiotemporal and LSTM networks (Figure 3D).

Importantly, the different models differ in the way they integrate spatial and temporal information along the hierarchy. These two types of information can be processed either sequentially, as is the case for the spatial-temporal network type that contains layers with one-dimensional filters that first integrate information across the different muscles, followed by an equal number of layers that integrate only in the temporal direction or simultaneously, using two-dimensional kernels, as they are in the spatiotemporal network. In the LSTM networks, spatial information was integrated similarly to the spatial-temporal networks, before entering the LSTM layer.

Candidate models for each class can be created by varying hyper-parameters such as the number of layers, number and size of spatial and temporal filters, type of regularization and response normalization, among others (see Table 2, Methods). As a first step to restrict the number of models, we performed a hyper-parameter architecture search by selecting models according to their performance on the proprioceptive tasks. We should emphasize that our ANNs are simultaneously integrating proprioceptive inputs and time, unlike standard feed-forward CNN-models of the visual pathway that just operate on images (23). TCNs have been shown to be excellent for time-series modeling (44), and therefore naturally describe neurons along a sensory pathway that integrate spatio-temporal inputs.

**Table 2.** Hyper-parameters for neural network architecture search. To form candidate networks, first a number of layers (per type) is chosen, ranging from 2-8 (in multiples of 2) for spatial-temporal models and 1-4 for the spatiotemporal and LSTM ones. Next, a spatial and temporal kernel size per Layer is picked where relevant, which remains unchanged throughout the network. For the spatiotemporal model, the kernel size is equal in both the spatial and temporal directions in each layer. Then, for each layer, an associated number of kernels/feature maps is chosen such that it never decreases along the hierarchy. Finally, a spatial and temporal stride is chosen. For the LSTM networks, the number of recurrent units is also chosen. All parameters are randomized independently and 50 models are sampled per network type. **Columns 2-4** Hyper-parameter values for the top-performing models in the ART. The values given under the “spatial” rows count for both the spatial and temporal directions for the spatiotemporal model.

|  | Hyper-parameters | Spatial-Temporal | Spatiotemporal | LSTM |
| --- | --- | --- | --- | --- |
| Num. Layers | [1, 2, 3, 4] | 4 + 4 | 4 | 3 + 1 |
| Spatial Kernels (pL) | [8, 16, 32, 64] | [8, 16, 16, 32] | [8, 8, 32, 64] | [8, 16, 16] |
| Temporal Kernels (pL) | [8, 16, 32, 64] | [32, 32, 64, 64] | n/a | n/a |
| Spatial Kernel Size | [3, 5, 7, 9] | 7 | 7 | 3 |
| Temporal Kernel Size | [3, 5, 7, 9] | 2 | n/a | n/a |
| Spatial Stride | [1, 2] | 9 | 2 | 1 |
| Temporal Stride | [1, 2, 3] | 3 | n/a | n/a |
| Num. Recurrent units | [128, 256] | n/a | n/a | 256 |

### Architecture search & representational changes

To find models that could solve the proprioceptive tasks, we performed an architecture search and trained 150 models (50 models per type). Notably, we trained the same model (as specified by architectural parameters) on both tasks by modifying the output and the loss function used to train the model. All networks were trained using the Adam optimizer (45) until performance on the validation set saturated. After training, all models were evaluated on a unseen test set (Figure S1A).

Models of all three types achieved excellent performance on the action recognition task (ART) (Figure 3E; multi-class accuracy of 98.86% ± 0.04, Mean ± SEM for the best spatial-temporal model, 97.93% ± 0.03 for the best spatiotemporal model, and 99.56% ± 0.04 for the best LSTM model, *N* = 5 randomly initialized models). The parameters of the best performing architectures are displayed in Table 2. The same models could also accurately solve the trajectory decoding task (TDT) (Figure 3E; with decoding errors of only 0.22 cm ± 0.005, Mean ± SEM for the best spatial-temporal model, .13 cm ± 0.003 for the best spatiotemporal model, and 0.05 cm ± 0.01 for the best LSTM model, *N* = 5 randomly initialized models). Of the hyper-parameters considered, the depth of the networks influenced performance the most (Figures 3E, S1B). Further, the performance on the two tasks were related: models performing well on one task tend to perform well on the other 3E.

Having found models that robustly solve the ART and TDT, we sought to analyze their properties. We created five pretraining (control) and post-training (trained) pairs of models for the best-performing model architecture for further analysis. We will refer to those as “instantiations”. As expected, the randomly initialized models performed at chance level (5%) on the ART.

How did the population activity change across the layers after learning the tasks? Initially, we focus on the best spatial-temporal model and then show that our analysis extends to the other model types. We compared the representations across different layers for each trained model to its random initialization by linear Centered Kernel Alignment (CKA, see Methods). This analysis revealed that for all instantiations the representations remained similar between the trained and control models for the first few layers and then deviate in the middle to final layers of the network (Figure 4A). Furthermore, trained models not only differed from the random initialization, but also across tasks, and the divergence appeared earlier (Figure 4A). Therefore, we found that both learning and the task substantially change the representations. Next, we aimed to understand how the tasks are solved, i.e., how the different stimuli are transformed across the hierarchy.

**Figure 4.**
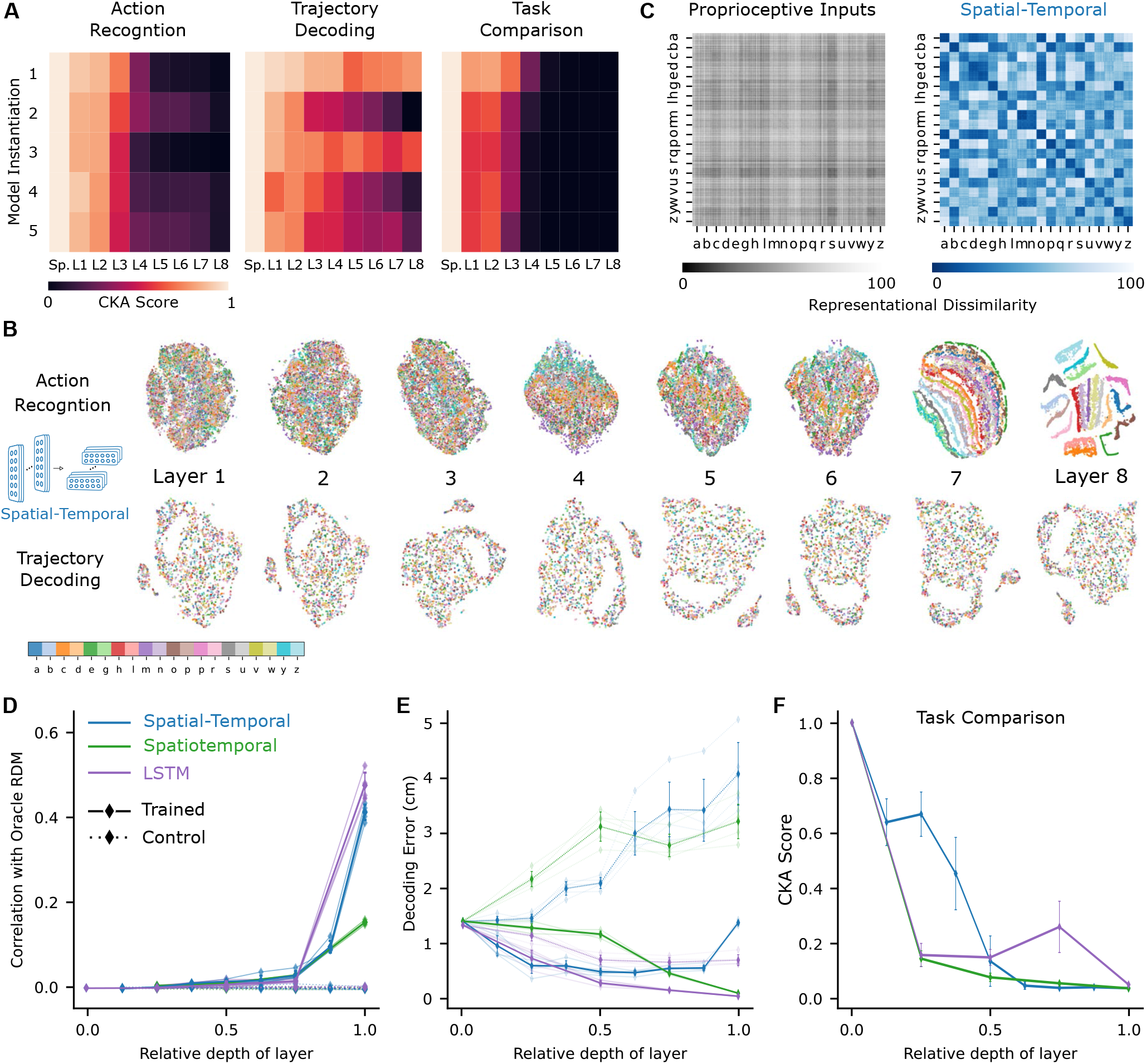
Low-dimensional embedding of network layers reveals structure. (**A**) Similarity in representations (CKA) between the trained and the control models for each of the five instantiations of the best performing spatial-temporal models (left and center). CKA between models trained on recognition vs decoding (right). (**B**) t-distributed stochastic neighbor embedding (t-SNE) for each layer of one instantiation of the best performing spatial-temporal model trained on both tasks. Each data point is a random stimulus sample (*N* = 2, 000, 50 per stimulus). (**C**) Representational Dissimilarity Matrices (RDM). Character level representation are calculated through percentile representational dissimilarity matrices for proprioceptive inputs (left) and final layer features (right) of one instantiation of the best performing spatio-temporal model trained on recognition task. (**D**) Similarity in stimulus representations between representational dissimilarity matrices of an Oracle (ideal observer) and each layer for the five instantiations of the ART-trained models and their controls. (**E**) Decoding error (in cm) along the hierarchy for each model type trained on the decoding task. (**F**) CKA between models trained on recognition vs decoding for the five instantiations of all network types (right).

To illustrate the geometry of the ANN representations and how the different characters are disentangled across the hierarchy we used t-SNE to visualize the structure. For the ART, the different characters remain entangled in the representations throughout most of the processing hierarchy before separating in the final layers (spatial-temporal model: Figure 4B and for the other model classes S2A). To quantify this, we computed representational dissimilarity matrices (RDM; see Methods). We found that different instances for the same characters were not represented similarly at the level of proprioceptive inputs, but at at the level of the last convolutional layer for the trained models (Figure 4C and for other model classes S2B). To quantify how the characters are represented across the hierarchy, we computed the similarity to an Oracle’s RDM, where an Oracle (or ideal observer) would have a block structure, with dissimilarity 0 for all stimuli of the same class and 1 (100th percentile) otherwise (Figure S2B). We found for all model instantiations similarity only increased towards the last layers (Figure 4D). This finding corroborates the visual impression gained via t-SNE that different characters are disentangled near the end of the processing hierarchy (Figures 4B, S2A, C).

How is the TDT solved across the hierarchy? In contrast to the ART trained models, as expected, representations of characters remained entangled throughout (Figure 4B, S2A). We found that the end-effector position can be decoded across the hierarchy (Figure 4E). This result is expected, as even from the proprioceptive input a linear readout achieves good performance (Figure 3C). Finally, we quantified CKA scores across the different architecture classes and found that with increasing depth the representations diverge between the two tasks (Figures 4F, S2C). Collectively, this suggests that characters are not immediately separable in ART-models, but the end-effector can be well decoded in TDT-models throughout the architecture.

### Single unit tuning properties

To gain insight into why ART- and TDT-trained models differ in their representations, we examined single unit tuning properties. In the primate these have been well described (3, 14), thus present as an ideal comparison point. Specifically, we analyzed the units for end-effector position, speed, direction, velocity, and acceleration tuning. We performed these analyses by relating variables (such as movement direction) to the activity of single units during the continuous movement (see Methods). Units with a test-*R*^2^ *>* 0.2 were considered “tuned” to that feature (this is a conservative value in comparison to experimental studies, e.g., 0.07 for (14)).

Given the precedence in the literature, we focused on direction tuning in *all horizontal* planes. We fit directional tuning curves to the units with respect to the instantaneous movement direction. As illustrated in examples, the ART spatial-temporal model (as well as proprioceptive inputs), directional tuning can be observed for the typical units shown (Figure 5A, B). Spindle afferents are known to be tuned to motion, i.e. velocity and direction (46). We verified the tuning of the spindles and found that the spindle component, tuned for muscle length, is primarily tuned for position (median *R*^2^ = 0.40, *N* = 25) rather than kinematics (median direction *R*^2^ = 0.0031, median velocity *R*^2^ = 0.0026, *N* = 25), whereas the spindle component tuned for changes in muscle length were primarily tuned for kinematics (median direction *R*^2^ = 0.55, velocity *R*^2^ = 0.81, *N* = 25), and only poorly tuned for position (median *R*^2^ = 0.34, *N* = 25). For the ART model, direction selectivity was prominent in middle layers 1-6 before decreasing by layer 8, and a fractions of units exhibited tuning to other kinematic variables with *R*^2^ *>* 0.2 (Figures 5C, S3A,B).

**Figure 5.**
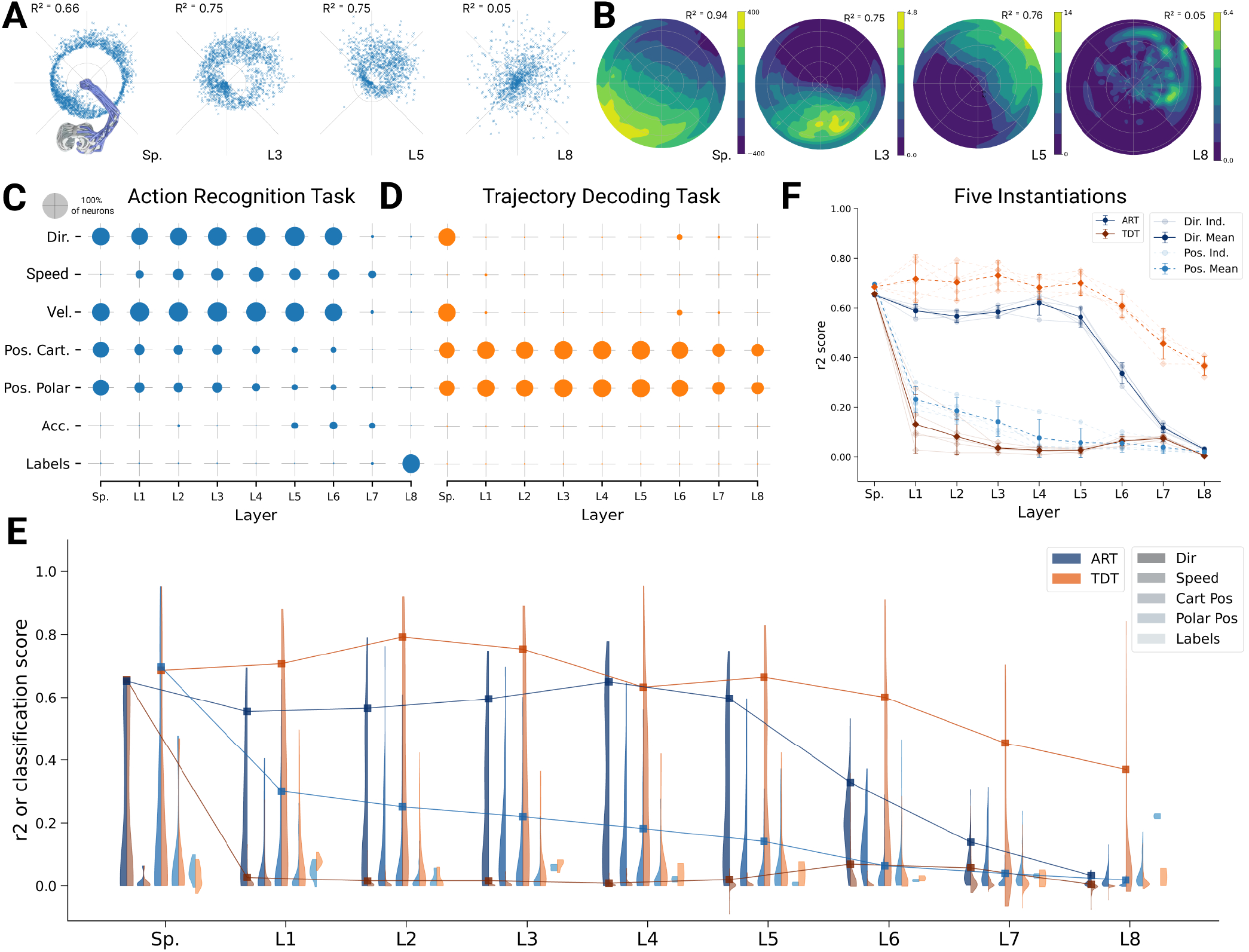
Analysis of single unit tuning properties for spatial-temporal models. (**A**) Polar scatter plots showing the activation of units (radius *r*) as a function of end-effector direction as represented by the angle *θ* for directionally tuned units in different layers of the top-performing spatial temporal model trained on the action recognition task. Directions correspond to that of the end-effector while tracing characters in the model workspace. The activation strengths of one (velocity-dependent) muscle spindle, one unit in layer 3, 5 and 8 of the are shown. (**B**) Similar to A, except that now radius describes velocity, and color represents activation strength. The contours are determined following linear interpolation, with gaps filled in by neighbor interpolation, and results smoothed using a Gaussian filter. Examples of one muscle spindle, one unit in layer 3, 5 and 8 are shown. (**C**) For each layer of one trained instantiation, the units are classified into types based on their tuning. A unit was classified as belonging to a particular type if its tuning had a test *R*^2^ *>* 0.2. Tested features were direction tuning, speed tuning, velocity tuning, Cartesian and polar position tuning, acceleration tuning, and label tuning (14/3890 scores excluded for ART-trained, 291/3890 for TDT-trained; see Methods). (**D**) The same plot but for the spatial-temporal model of the same architecture but trained on the trajectory decoding task. (**E**) For an example instantiation, the distribution of test *R*^2^ scores for both the ART and TDT trained models are shown, for five kinds of kinematic tuning for each layer: direction tuning, speed tuning, Cartesian position tuning, polar position tuning, and label-specificity indicated by different shades and arranged left-right for each layer including spindles. The solid line connects the 90%-quantiles of two of the tuning curve types, direction tuning (dark) and position tuning (light). Tuning scores were excluded if they were equal to 1, indicating a constant neuron, or less than − 0.1, indicating an improper fit (8 scores excluded; see Methods). (**F**) The means of 90%-quantiles over all five model instantiations of models trained on ART and TDT are shown for direction tuning (dark) and position tuning (light). 95%-confidence intervals are shown over instantiations (*N* = 5).

In contrast, for the TDT model, no directional tuning was observed, but positional tuning (Figure 5D). These observations are further corroborated when comparing the distributions of tuning properties (Figure 5E) and 90%-quantiles for all the instantiations (Figure 5F). The difference in median tuning score between the two differently trained groups of models across the five model instantiations becomes significant starting in the first layer for both direction (layer 1 *t*(4) = 13.07, p=0.0002; layer 2 *t*(4) = 15.79, p=0.0001; layer 3 *t*(4) = 39.74, p=2.65e-06; layer 4 *t*(4) = 5.24, p=5.24e-06; layer 5 *t*(4) = 35.76, p=3.65e-06; layer 6 *t*(4) = 12.97, p=0.0002; layer 7 *t*(4) = 3.605, p=0.017; *t*(4) = 19.35, p=4.21e-05) and position (layer 1 *t*(4) = −11.21, p=0.0004; layer 2 *t*(4) = −28.59, p=8.91e-06; layer 3 *t*(4) = −21.27, p=2.89e-05; layer 4 *t*(4) = −13.91, p=0.0002; layer 5 *t*(4) = −18.55, p=4.97e-05; layer 6 *t*(4) = −61.59, p=4.16e-07; layer 7 *t*(4) = −24.08, p=1.77e-05; *t*(4) = −25.80, p=1.34e-05).

Given that the ART models are trained to recognize characters, we asked if single units are well tuned for specific characters. To test this we trained an SVM to classify characters from the single unit activations. Even in the final layer (before the readout) of the spatial-temporal model, the median classification performance over the five model instantiations as measured by the normalized area under the ROC curve-based selectivity index for single units was 0.212 ± 0.005 (mean ± SEM, *N* = 5 instantiations), and was never higher than 0.41 for any individual unit across all model instantiations (see Methods). Thus, even in the final layer there are effectively no single-character specific units. Of course, combining the different units of the final fully connected layer gives a high fidelity readout of the character and allows the model to achieve high classification accuracy. Thus, character identity is represented in a distributed way. In contrast, and as expected, character identity is poorly encoded in single cells for the TDT model (Figure 5D,F).

These main results also hold for the other architecture classes. In spatiotemporal models, in which both spatial and temporal processing occurs between all layers, we observe a monotonic decrease in the directional tuning across the four layers for the ART task and a quick decay for the TDT task (Figure 6 A, B). Speed and acceleration tuning are present in the ART, but not in the TDT models (Figure S3B, C). Conversely, we find that positional coding is stable for TDT models and not the ART models. The same results hold true for LSTM model (Figures 6 C, D, S3E, F). The differences in directional and Cartesian positional tuning were statistically significant for all layers according to a paired t-test with 4 degrees of freedom for both model types. Thus, for all architecture classes we find that strong direction selective tuning is present in early layers of models trained with the ART task, but not the TDT task.

**Figure 6.**
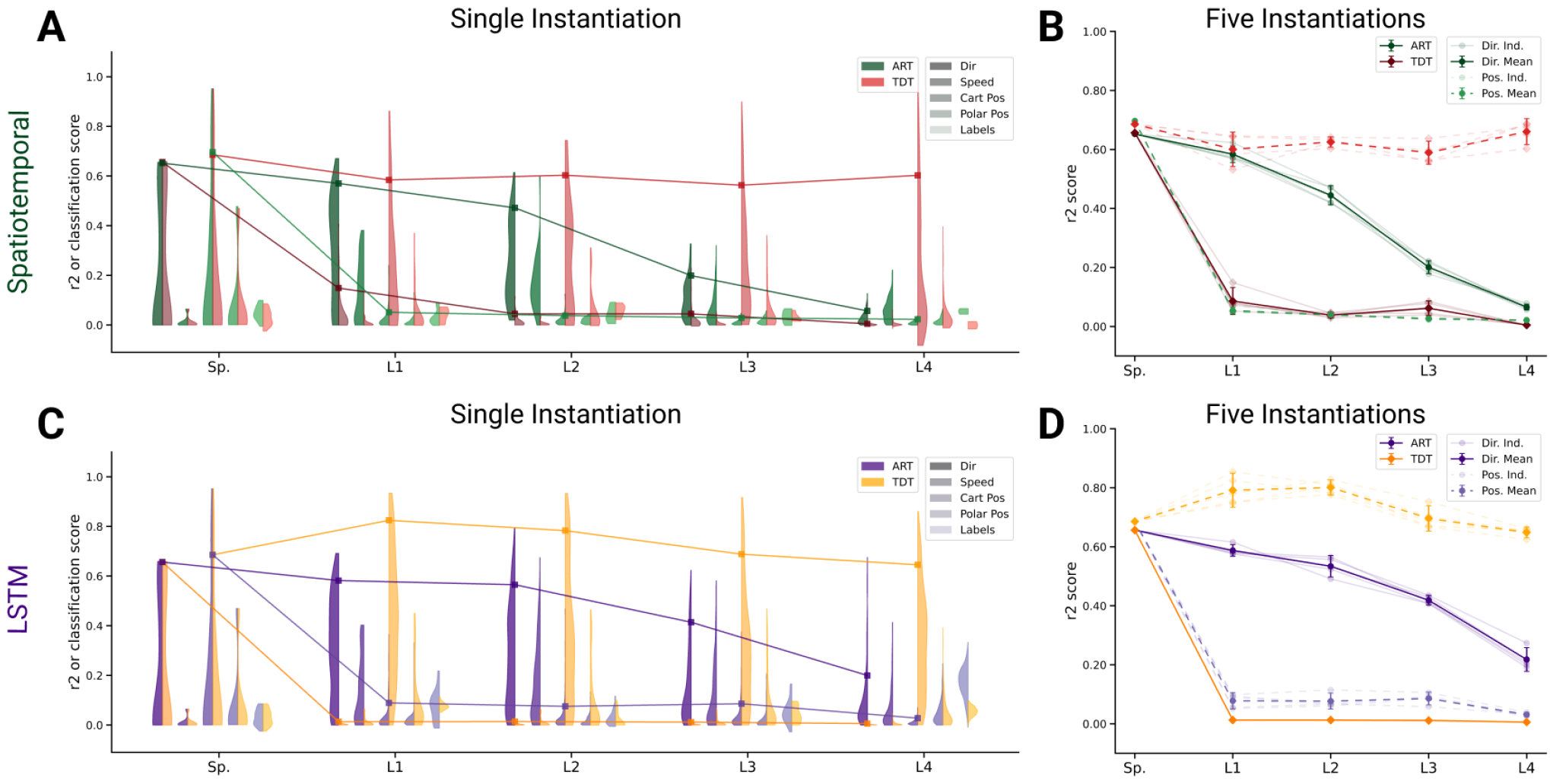
Analysis of single unit tuning properties for spatiotemporal and LSTM models. (**A**) For an example instantiation of the top-performing spatiotemporal model, the distribution of test *R*^2^ scores for both the trained and control model are shown, for five kinds of kinematic tuning for each layer: direction tuning, speed tuning, Cartesian position tuning, polar position tuning, and label-specificity. The solid line connects the 90%-quantiles of two of the tuning curve types, direction tuning (dark) and position tuning (light). (0/2330 scores excluded summed over all layers for ART-trained, 129/2330 for TDT; see Methods). (**B**) The means of 90%-quantiles over all five model instantiations of models trained on action recognition and trajectory decoding are shown for direction tuning (dark) and position tuning (light). 95%-confidence intervals are shown over instantiations (*N* = 5). (**C**) The same plot as in (A) but for the top-performing LSTM model (0/6530 scores excluded for ART-trained, 1024/6530 for TDT; see Methods). (**D**) The same plot as B, for the LSTM model.

Our results so far suggests that the primate proprioceptive pathway is more consistent with the action recognition hypothesis, but to corroborate this, we also assessed decoding performance, which measures representational information. For all architecture types movement direction and speed can be better decoded from ART than from TDT trained models (Figure S4A,C,E). In contrast, for all architectures, position can be better decoded for TDT than for ART trained models (Figure S4B,D,F). These results are consistent with the single-cell encoding results, and again lend support for the proprioceptive systems involvement in action representation.

So far, we have directly compared TDT and ART models. This does not address task-training as such. Namely, we found directional selective units in ART-models and positional-selective units in TDT-models, but how do those models compare to randomly initialized models (controls)? Remarkably, directional selectivity is similar for ART and control models (Figure S5). In contrast to controls, ART-trained models harbor less positional tuned units. The situation is reversed for TDT-trained models – those models gain positionally tuned units and lose directionally selective units during task training (Figure S6). Consistent with those encoding results, position can be worse and direction and speed similarly decoded from ART-models than from controls, respectively (Figure S7). Conversely, direction and speed can be worse and position better decoded from TDT-models than controls (Figure S7).

### Uniformity and coding invariance

We compared population coding properties to further elucidate the similarity to S1. We measured the distributions of preferred directions and whether coding properties are invariant across different workspaces (reaching planes). Prud’homme and Kalaska found a relatively uniform distribution of preferred directions in primate S1 during a center out reaching 2D manipulandum-based task (Figure 7A from (14)). In contrast, most velocity tuned spindle afferents have preferred directions located along one major axis pointing frontally and slightly away from the body (Figure 7B). Qualitatively, it appears that the ART trained model had more uniformly distributed preferred directions in the middle layers compared to control models with randomly initialized weights (Figure 7C).

**Figure 7.**
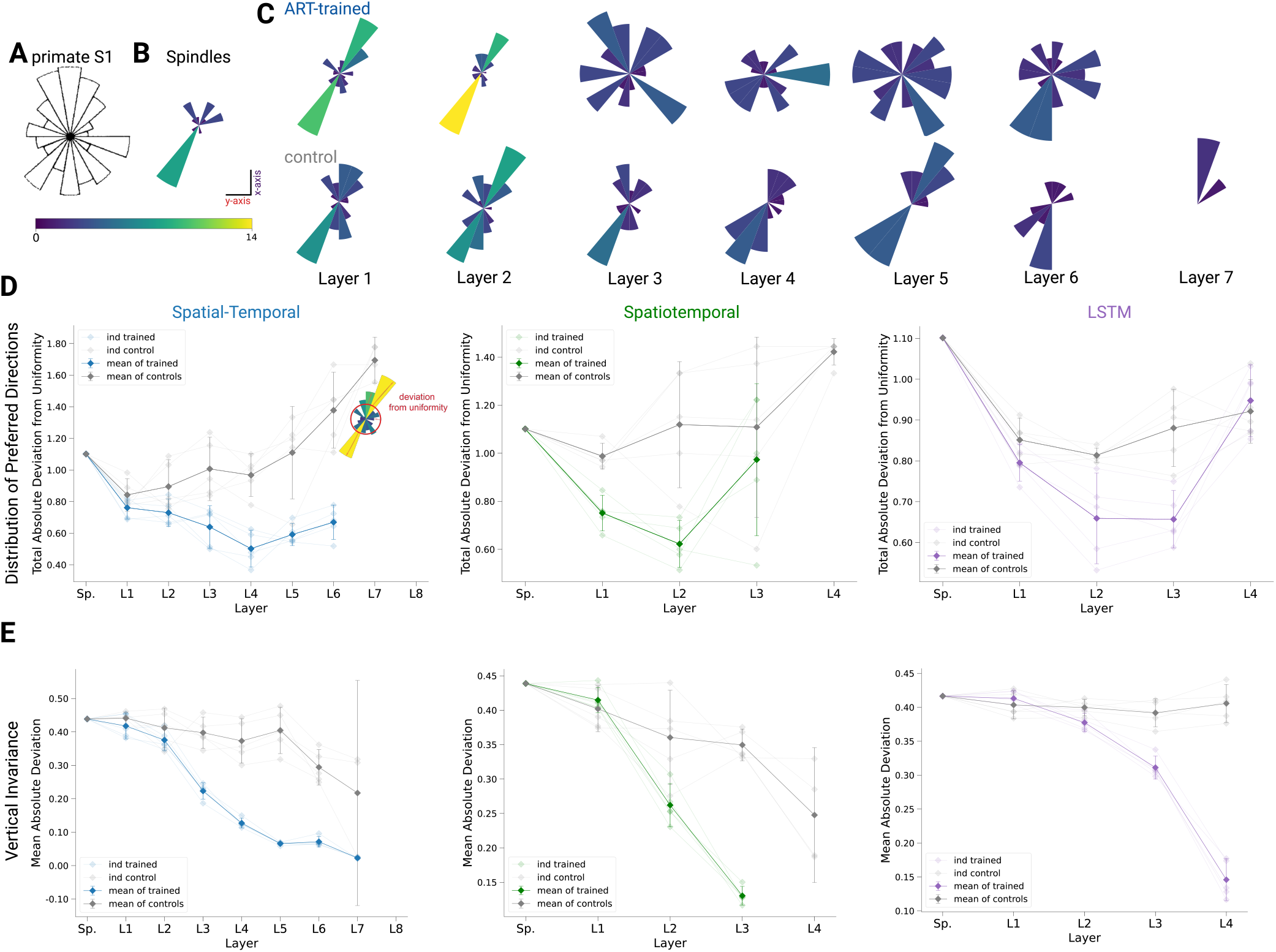
Distribution of preferred directions and invariance of representation across workspaces. (**A**) Adopted from (14); distribution of preferred directions in primate S1. (**B**) Distribution of preferred directions for spindle input. (**C**) Distribution of preferred directions for one spatial-temporal model instantiation (all units with *R*^2^ *>* 0.2 are included). Bottom: the corresponding control. For visibility all histograms are scaled to the same size and the colors indicate the number of tuned neurons. (**D**) For quantifying uniformity, we calculated the total absolute deviation from the corresponding uniform distribution over the bins in the histogram (red line in inset) for the spatial-temporal model (*left*), the spatiotemporal model (*middle*), and the LSTM model (*right*). Normalized absolute deviation from uniform distribution for preferred directions per instantiation are shown (*N* = 5, faint lines) for trained and control models as well as mean and 95%-confidence intervals over instantiations (solid line; *N* = 5). Note that there is no data for layers 7 and 8 of the trained spatial-temporal model, layer 8 of the untrained spatial-temporal model, and layer 4 of the spatiotemporal model as they have no direction selective units (*R*^2^ *>* 0.2). (**E**) For quantifying invariance we calculated mean absolute deviation in preferred orientation for units from the central plane to each other vertical plane (for units with *R*^2^ *>* 0.2). Results are shown for each instantiation (*N* = 5, faint lines) for trained and control models plus mean (solid) and 95%-confidence intervals over instantiations (*N* = 5). Note that there is no data for layer 4 of the trained spatiotemporal model, as it have no direction selective units (*R*^2^ *>* 0.2).

To quantify uniformity we calculated the total absolute deviation (TAD) from uniformity in the distribution of preferred directions. The results indicate that the distribution of preferred directions becomes more uniform in middle layers for all instantiations of the different model architectures (Figure 7D), and that this difference is statistically significant for the spatial-temporal model beginning in layer 3 (layer 3 *t*(4) = − 3.41, p=0.027; layer 4 *t*(4) = − 8.12, p=0.001; layer 5 *t*(4) = − 4.08, p=0.015; layer 6 *t*(4) = − 5.55, p=0.005). This analysis revealed that while randomly initialized models also have directionally selective units, those units are less uniformly distributed than in models trained with the ART task. Importantly, TDT trained modules have almost no directionally selective units, further corroborating that ART models are *more consistent* with Prud’homme and Kalaska’s findings (14).

Lastly, we could directly test if preferred tuning directions (of tuned units) were maintained across different planes due to the fact that we created trajectories in multiple vertical and horizontal planes. We hypothesized that for the trained networks preferred orientations would be more preserved across planes compared to controls. In order to examine how an individual unit’s preferred direction changed across different planes, directional tuning curve models were fit in each horizontal/vertical plane separately (examples in Figure S9A, B). To measure the representational invariance, we took the mean absolute deviation (MAD) of the preferred tuning direction for directionally tuned units (*R*^2^ *>* 0.2) across planes (see Methods) and averaged over all planes (Figures 7E, S9A). For the spatial-temporal model across vertical workspaces, layers 3-6 were indeed more invariant in their preferred directions (layer 3: t(4)= -11.38, p=0.0003; layer 4: t(4) = -10.23, p=0.0005; layer 5: t(4) = -12.17, p=0.002; layer 6: t(4) = - 13.18, p=0.0002; Figure 7E; variation in preferred direction illustrated for layer 5 in Figure S9D for trained model and in Figure S9E for controls). The difference in invariance for the horizontal planes was likewise statistically significant for layers 4-6 (Figure S9A). The difference in invariance here might only become statistically significant one layer later because the spindles are already more invariant in the horizontal planes (MAD: 0.225 ± 2.78*e* − 17, *mean* ± *SEM*, N = 25; Figure S9B) than the vertical workspaces (MAD: 0.439 ± 5*e* − 17, mean ± SEM, N = 16; Figure S9C), meaning that it takes a greater amount of invariance in the trained networks for differences with the control networks to become statistically apparent. For the spatiotemporal and LSTM models, the relatively slower increase in invariance in the horizontal direction is exaggerated even more. For the LSTM model, the neuron tuning does not become stronger for the horizontal planes until the recurrent laye. The spatiotemporal trained models even begin by being less invariant in the horizontal planes before the difference evens out in layer 3r (FigureS9A).

## Discussion

### Task-Driven Modeling of Proprioception

For various anatomical and experimental reasons, recording proprioceptive activity during natural movements is technically challenging (3, 47). Furthermore, “presenting” particular proprioceptive-only stimuli is difficult, which poses substantial challenges for systems identification approaches. This highlights the importance of developing accurate, normative models that can explain neural representations across the proprioceptive pathway, as has been successfully done in the visual system (20–26). To tackle this, we combined human movement data, biomechanical modeling, as well as deep learning to provide a blueprint for studying the proprioceptive pathway.

We presented a task-driven approach to study the proprioceptive system based on our hypothesis that proprioception can be understood normatively as having to solve action-recognition from receptor inputs. We created a passive character recognition task for a simulated human biomechanical arm paired with a muscle spindle model and found that deep neural networks can be trained to accurately solve the action recognition task. Inferring the character from passive arm traces was chosen as it is a type of task that humans can easily perform and because it covers a wide range of natural movements of the arm. The perception is also likely fast, so that feed-forward processing is a good approximation (while we also find similar results with recurrent models). Additionally, character recognition is an influential task for studying ANNs, for instance MNIST (42, 48). Moreover, when writing movements were imposed onto the ankle with a fixed knee joint, the movement trajectory could be decoded from a few spindles using a population vector model, suggesting that spindle information is accurate enough for decoding (49). Lastly, while the underlying movements are natural and of ethological importance for humans, the task itself is only a small subset of human upper-limb function. Thus, it posed an interesting question whether such a task would be *sufficient* to induce representations similar to biological neurons.

We put forth a normative model of the proprioceptive system, which is experimentally testable. This builds on our earlier preprint (50) of this work, which only considered action recognition and used a different receptor model (51) that used changes in the length of the muscles. Here, we also include positional sensing and an additional tasks to directly test our hypothesis of action coding vs. the canonical view of proprioception. We confirm that in ART-trained models, but not in randomly initialized models, the intermediate representations contain directionally selective neurons that are uniformly distributed (50). Furthermore, we had predicted that earlier layers and in particular muscle spindles, have a biased, bidirectionally tuned distribution. This distribution was later found experimentally for single units in the cuneate nucleus (52). Here we still, robustly find this result but with different spindle models (35, 36). However, these PD distribution result does not hold when the identical architectures are trained with the trajectory decoding task. In those models directionally tuned neurons do not emerge, in fact they are “unlearned” in comparison to random initializations for TDT (Figure S6).

The distribution of preferred directions becomes more uniform over the course of the processing hierarchy (Figure 6 and 7), similar to the distribution of preferred tuning in somatosensory cortex (14). This does not occur in the randomized controls, which instead maintained an input distribution centered on the primary axis of preferred directions of the muscular tuning curves. Furthermore, the task-trained models make a prediction about the distribution of preferred directions along the proprioceptive pathway. For instance, we predict that in the brainstem - i.e. cuneate nucleus - preferred directions are aligned along major axes inherited from muscle spindles that correspond to biomechanical constraints (consistant with Versteeg et al. (52)). A key element of robust object recognition is invariance to task-irrelevant variables (23, 53). In our computational study, we could probe many different workspaces (26 horizontal and 18 vertical) to reveal that training on the character recognition task makes directional tuning more invariant (Figure 7E). This together with our observation that directional tuning is simply inherited from muscle spindles, highlights the importance of sampling the movement space well, as also emphasized by pioneering experimental studies (54). We also note that the predictions depend on the musculoskelatal model and the movement statistics. In fact, we predict that e.g., distributions of tuning and invariances might be different in mice, a species that has a different body orientation from primates. This will be explored in future studies.

### Limitations and future directions

Our model only encompasses proprioception and trained in a supervised fashion. However, it is quite natural to interpret the supervised feedback stemming from other senses. For instance, the visual system could naturally provide information of the hand localization or about the type of character.

Here we used different types of temporal convolutional and recurrent network architectures. In future work it will be important to investigate emerging, perhaps more biologically-relevant architectures to better understand how muscle spindles are integrated in upstream circuits. While we used spindle Ia and II models, it is known that multiple receptors, namely cutaneous, joint, and muscle receptors play a role for limb localization and kinesthesia (3, 55–58). For instance, a recent simulation study by Kibleur et al. highlighted the complex spatio-temporal structure of proprioceptive information at the level of the cervical spinal cord (47). Furthermore, due to fusimotor drive receptor activity can be modulated by other modalities, e.g., vision (59). In the future, models for other afferents, golgi tendon organ incl. the fusimotor drive as well as cutaneous receptors can be added, to study their role in the context of various tasks (40).

### Conclusions

We proposed action recognition from peripheral inputs as an objective to study proprioception. We developed task-driven models of the proprioceptive system and showed that diverse preferred tuning directions and invariance across 3D space emerges in neural networks emerges in such systems – this emergence was not found for models trained on representing the state of the body. Due to their hierarchical nature, the network models provide not only a description of neurons previously found in physiology studies, they make predictions about coding properties, such as the biased distribution of direction tuning in subcortical areas.

## Methods

### Proprioceptive Character Trajectories: Dataset and Tasks

#### The character trajectories dataset

The movement data for our task was obtained from the UCI Machine Learning Repository character trajectories dataset (33, 37). In brief, the dataset contains 2,858 pen-tip trajectories for 20 single-stroke characters (excluding f, i, j, k, t and x, which were multi-stroke in this dataset) in the Latin alphabet, written by a single person on an Intuos 3 Wacom digitization tablet providing pen-tip position and pressure information at 200 Hz. The size of the characters was such that they all approximately fit within a 1 × 1 cm grid. Since we aimed to study the proprioception of the whole arm, we first interpolated the trajectories to lie within a 10 × 10 cm grid and discarded the pen-tip pressure information. Trajectories were interpolated linearly while maintaining the velocity profiles of the original trajectories. Empirically, we found that on average it takes 3 times longer to write a character in the 10 × 10 cm grid than in the small 1 × 1 one. Therefore, the time interval between samples was increased from 5ms (200 Hz) to 15ms (66.7 Hz) when interpolating trajectories. The resulting 2,858 character trajectories served as the basis for our end-effector trajectories.

#### Computing joint angles and muscle length trajectories

Using these end-effector trajectories, we sought to generate realistic proprioceptive inputs while passively executing such movements. For this purpose, we used an open-source musculoskeletal model of the human upper limb, the upper extremity dynamic model by Saul et al. (34, 60). The model includes 50 Hill-type muscle-tendon actuators crossing the shoulder, elbow, forearm and wrist. While the kinematic foundations of the model enable it with 15 degrees of freedom (DoF), 8 DoF were eliminated by enforcing the hand to form a grip posture. We further eliminated 3 DoF by disabling the model to have elbow rotation, wrist flexion and rotation. The four remaining DoF are elbow flexion (*θ*_*ef*_), shoulder rotation (*θ*_*sr*_), shoulder elevation i.e, thoracohumeral angle (*θ*_*se*_) and elevation plane of the shoulder (*θ*_*sep*_).

The first step in extracting the spindle activations involved computing the joint angles for the 4 DoF from the endeffector trajectories using constrained inverse kinematics. We built a 2-link 4 DoF arm with arm-lengths corresponding to those of the upper extremity dynamic model (60). To determine the joint-angle trajectories, we first define the forward kinematics equations that convert a given joint-angle configuration of the arm to its end-effector position. For a given joint-angle configuration of the arm **q** = [*θ*_*ef*_, *θ*_*sr*_, *θ*_*se*_, *θ*_*sep*_]^*T*^, the end-effector position **e** ∈ ℝ^3^ in an absolute frame of reference {*S*} centered on the shoulder is given by

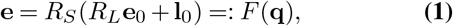

with position of the end-effector (hand) **e**_0_ and elbow **l**_**0**_ when the arm is at rest and rotation matrices

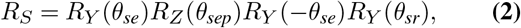

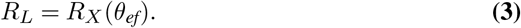

Thereby, *R*_*S*_ is the rotation matrix at the shoulder joint, *R*_*L*_ is the rotation matrix at the elbow obtained by combinations of intrinsic rotations around the X, Y and Z axes which are defined according to the upper extremity dynamic model (60), treating the joint angles as Euler angles and *R*_*X*_, *R*_*Y*_, *R*_*Z*_ - the three basic rotation matrices.

Given the forward kinematics equations, the joint-angles **q** for an end-effector position **e** can be obtained by iteratively solving a constrained inverse kinematics problem for all times *t* = 0 … *T* :

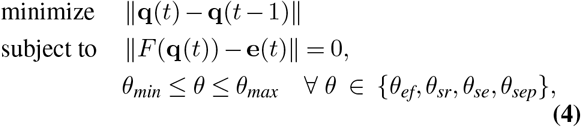

Where **q**(− 1) is a natural pose in the center of the workspace (see Figure 2D) and each **q**(*t*) is a posture pointing to **e**(*t*), while being close to the previous posture **q**(*t* − 1). Thereby, {*θ*_*min*_, *θ*_*max*_} define the limits for each joint-angle. For a given end-effector trajectory **e**(*t*), joint-angle trajectories are thus computed from the previous time point in order to generate smooth movements in joint space. This approach is inspired by D’Souza et al. (61).

Finally, for a given joint trajectory **q**(*t*) we passively moved the arm through the joint-angle trajectories in the OpenSim 3.3 simulation environment (62, 63), computing at each time point the equilibrium muscle lengths **m**(*t*) ∈ ℝ^25^, since the actuation of the 4 DoFs is achieved by 25 muscles. For simplicity, we computed equilibrium muscle configurations given joint angles as an approximation to passive movement.

#### Proprioceptive inputs

While several mechanoreceptors provide proprioceptive information, including joint receptors, Golgi tendon organs and skin stretch receptors, the muscle spindles are regarded as the most important for conveying position and movement related information (2, 51, 64, 65). Here we are inspired by Dimitriou and Edin’s recordings from human spindles (35, 36). They found that both Ia and II units are well predicted by combinations (for parameters *k*_1_ … *k*_5_) of muscle length *l*, muscle velocity *l*′, acceleration *l*″ and EMG:

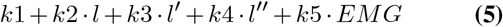

As we model passive movement, the associated EMG activity is negligible. To simplify the aggregate information flowing from one muscle (via multiple Ia and II spindles), we consider a more generic/functional representation of proprioceptive information as consisting of muscle length and velocity signals, which are approximately conveyed by muscle spindles during passive movements. Therefore, in addition to the equilibrium muscle lengths **m**(*t*), we input muscle velocity **v**(*t*) signal obtained by taking the first derivative. Taken together, {**m**(*t*), **v**(*t*)} form the proprioceptive inputs to train models of the proprioceptive system.

#### A scalable proprioceptive character trajectories dataset

We move our arms in various configurations and write at varying speeds. Thus, several axes of variation were added to each (original) trajectory by (1) applying affine transformations such as scaling, rotation and shear, (2) modifying the speed at which the character is written, (3) writing the character at several different locations (chosen from a grid of candidate starting points) in the 3D workspace of the arm, and (4) writing the characters on either transverse (horizontal) or frontal (vertical) planes of which there were 26 and 18 respectively, placed at a spatial distance of 3cm from each other (see Table 1 for parameter ranges). We first generated a dataset of end-effector trajectories of 1 million samples by generating variants of each original trajectory, by scaling, rotating, shearing, translating and varying its speed. For each end-effector trajectory, we compute the joint-angle trajectory by performing inverse kinematics. Subsequently, we simulate the muscle length and muscle velocity trajectories. Since different characters take different amount of time to be written, we pad the movements with static postures corresponding to the starting and ending postures of the movement, and jitter the beginning of the writing to maintain ambiguity about when the writing begins.

From this dataset of trajectories, we selected a subset of trajectories such that the integral of joint-space jerk (third derivative of movement) was less than 1 rad/s^3^ so as to ensure that the arm movement is sufficiently smooth. Among these, we picked the trajectories for which the integral of muscle-space jerk was minimal, while making sure that the dataset is balanced in terms of the number of examples per class, resulting in 200,000 samples. The final dataset consists of muscle length and velocity trajectories from each of the 25 muscles over a period of 320 time points, simulated at 66.7 Hz (i.e, 4.8 seconds). In other words, the dimensionality of the proprioceptive inputs in our tasks is 25×320×2. The dataset was then split into a training, validation and test set with a 72 − 8 − 20 ratio.

#### Action recognition and trajectory decoding tasks

Having simulated a large scale dataset of proprioceptive character trajectories, we designed two tasks - (1) the action recognition task (ART) to classify the identity of the character based on the proprioceptive inputs, and (2) the trajectory decoding task (TDT) to decode the endeffector coordinates (at each time step) from proprioceptive inputs. Baseline models (SVMs for the ART and linear regression for TDT) were first trained to investigate the difficulty of the task, followed by a suite of deep neural networks that aim to model the proprioceptive pathway.

#### Low dimensional embedding of population activity

−

To visualize population activity (of kinematic or network representations), we created low-dimensional embeddings of the proprioceptive inputs (Figure 3A) as well as the the layers of the neural network models, along time, and space/muscles dimensions (Figure 4B and Figure S2A). To this end, we first used Principal Components Analysis (PCA) to reduce the space to 50 dimensions, typically retaining around 75−80% of the variance. We then used t-distributed stochastic neighbor embedding (t-SNE, 38) using sklearn (66) with a perplexity of 40 for 300 iterations, to reduce these 50 dimensions down to two for visualization.

#### Support vector machine (SVM) analysis for action recognition

To establish a baseline performance for multi-class recognition we used pairwise SVMs with the one-against-one method (39). That is, we train 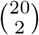 pairwise (linear) SVM classifiers (Figure 3C) and at test time implement a voting strategy based on the confidences of each classifier to determine the class identity. We trained SVMs for each input modality (end-effector trajectories, joint angle trajectories, muscle fiber-length trajectories and proprioceptive inputs) to determine how the format affects performance. All pairwise classifiers were trained using a hinge loss, and crossvalidation was performed with 9 regularization constants logarithmically spaced between 10^−4^ and 10^4^.

#### Baseline linear regression model for trajectory decoding

To establish how well one could decode endeffector coordinates from the joint, muscle and proprioceptive inputs, we trained linear regressors with ordinary least-squares loss using stochastic gradient descent until the validation loss saturated (with a tolerance of 10^−3^). Inputs and outputs to the model were first transformed using a standard scaler to center (remove the mean) and scale to unit variance over each feature in order to train faster. At test time, the same scalers were reused. Decoding error was determined as the squared error (L-2 norm) of the predicted and true endeffector coordinates in 3D.

### Models of the proprioceptive system

We trained two types of convolutional networks and one type of recurrent network on the two tasks. Each model is characterized by the layers used – convolutional and/or recurrent – which specify how the spatial and temporal information in the proprioceptive inputs is processed and integrated.

Each convolutional layer contains a set of convolutional filters of a given kernel size and stride, along with response normalization and a point-wise non-linearity. The convolutional filters can either be 1-dimensional, processing only spatial or temporal information or 2-dimensional, processing both types of information simultaneously. For response normalization we use layer normalization (67), a commonly used normalization scheme to train deep neural networks, where the response of a neuron is normalized by the response of all neurons of that layer. As point-wise non-linearity we use rectified linear units. Each recurrent layer contains a single LSTM (long short-term memory) cell with a given number of units that process the input one time step at a time.

Depending on what type of convolutional layers are used and how they are arranged, we classify convolutional models into two subtypes (1) spatial-temporal and (2) spatiotemporal networks. Spatial-temporal networks are formed by combining multiple 1-dimensional spatial and temporal convolutional layers. That is, the proprioceptive inputs from different muscles are first combined to attain a condensed representation of the ‘spatial’ information in the inputs, through a hierarchy of spatial convolutional layers. This hierarchical arrangement of the layers leads to increasingly larger receptive fields in spatial (or temporal) dimension that typically (for most parameters) gives rise to a representation of the whole arm at some point in the hierarchy. The temporal information is then integrated using temporal convolutional layers. In the spatiotemporal networks, multiple 2-d convolutional layers where convolutional filters are applied simultaneously across spatial and temporal dimensions are stacked together. The LSTM models on the other hand are formed by combining multiple 1-dimensional spatial convolutional layers and a single LSTM layer at the end of a stack of spatial filters that recurrently processes the temporal information. For each network, the features at the final layer are mapped by a single fully connected layer either onto a 20 dimensional (logits) or a 3 dimensional output (end-effector coordinates).

For each specific network type the following hyper-parameters were used: number of layers, number and size of spatial and temporal filters and their corresponding stride (see Table 2). Using this set of architectural parameters, 50 models of each type were randomly generated. Notably, we trained the same model (as specified by the architecture) on both tasks.

#### Network training and evaluation procedure

The action-recognition trained models were trained by minimizing the softmax cross entropy loss using the Adam Optimizer (45) with an initial learning rate of 0.0005, batch size of 256 and decay parameters (*β*_1_ and *β*_2_) of 0.9 and 0.999. During training the performance was monitored on the left-out validation set. When the validation error did not improve for 5 consecutive epochs, we decreased the learning rate by a factor of 4. After the second time the validation error saturated, we ended the training and evaluated accuracy of the networks on the test set. Overall we observe that the trained networks generalized well to the test data, even though the shallower networks tended to overfit S1A.

The TDT trained models on the other hand were trained to minimize the mean squared error between predicted and true trajectories. Hyperparameter settings for the optimizer, batch size and early stopping procedure used during training remained same across both tasks. Here, we observe that train and test decoding errors were highly correlated, and thereby achieve excellent generalization to test data S1A.

#### Comparison with controls

For each of the three types of models, the architecture belonging to the best performing model on the ART (as identified via the hyper-parameter search) was chosen as the basis of the analysis (Table 2). The resulting sizes of each layer’s representation across the hierarchy are given in Table 3. For each different model type, five sets of random weights were initialized and saved. Then, each instantiation was trained on both ART and TDT using the same training procedure as described in the previous section, and the weights were saved again after training. This gives a before and after structure for each run that allows us to isolate the effect of task-training.

**Table 3.**
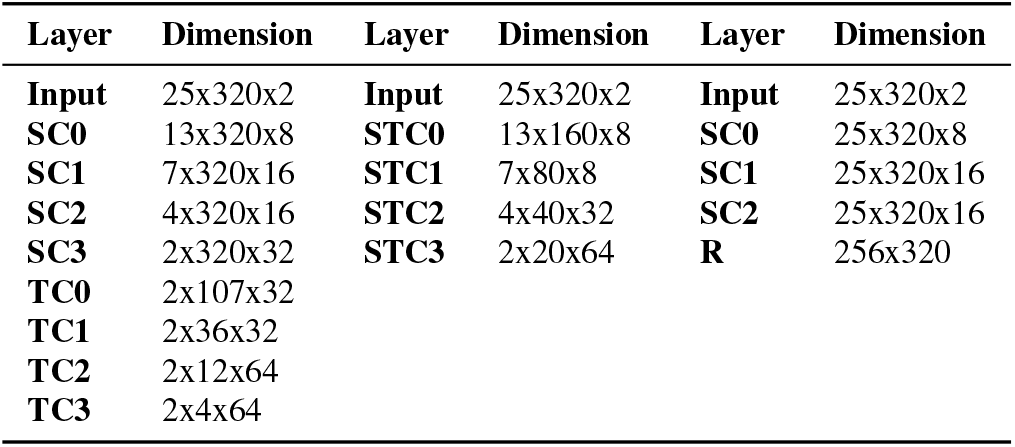
Size of representation at each layer for best performing architecture of each network type (spatial x temporal x filter dimensions).

### Population comparisons

#### Centered Kernel Analysis

In order to provide a population-level comparison between the trained and control models (Figure 4A), we used Linear Centered Kernel Analysis (CKA) for a high-level comparison of each layers’ activation patterns (68). CKA is an alternative that extends Canonical Correlation Analysis (CCA) by weighting activation patterns by the eigenvalues of the corresponding eigenvectors (68). As such, it maintains CCA’s invariance to orthogonal transformations and isotropic scaling, yet retains a greater sensitivity to similarities. Using this analysis, we quantified the similarity of the activation of each layer of the trained models with those of the respective controls in response to identical stimuli comprising 50% of the test set for each of the five model instantiations.

#### Representational similarity analysis

Representational Similarity Analysis (RSA) is a tool to investigate population level representations among competing models (69). The basic building block of RSA is a representational dissimilarity matrix (RDM). Given stimuli {*s*_1_, *s*_2_, …, *s*_*n*_} and vectors of population responses {*r*_1_, *r*_2_, …, *r*_*n*_}, the RDM is defined as:

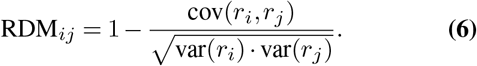

One of the main advantages of RDMs is that it characterizes the geometry of stimulus representation in a way that is independent of the dimensionality of the feature representations, so we can easily compare between arbitrary representations of a given stimulus set. Example RDMs for proprioceptive inputs, as well as the final layer before the readout for the best models of each type are shown in Figure 4C, S2B. Each RDM is computed for a random sample of 4,000 character trajectories (200 from each class) by using the correlation distance between corresponding feature representations. To compactly summarize how well a network disentangles the stimuli we compare the RDM of each layer to the RDM of the ideal observer, which has a RDM with perfect block structure (with dissimilarity values 0 for all stimuli of the same class and 1 (100 percentile) otherwise; see Figure S2B).

### Single unit analysis

#### Comparing the tuning curves

To elucidate the emerging coding properties of single units, we determined label specificity and fit tuning curves. Specifically, we focused on kinematic properties such as direction, velocity, acceleration and position of the end-effector for movements (Figure 5, 7). For computational tractability, 20, 000 of the original trajectories were randomly selected for the ART-trained S and ST models, and 4000 for TDT-trained ones as well as for both kinds of LSTM models. Each time point was treated as its own independent sample. In convolutional layers in which the hidden layers had a reduced temporal dimensionality than the input, the input trace was downsampled. Only those time points were kept that correpond to the center of the receptive fields of the units in the hidden layers.

A train-test split of 80 − 20 was used. The tuning curves were fit and tested jointly on all movements in planes with a common orientation, vertical or horizontal. The analysis was repeated for each of the five trained and control models. For each of the five different types of tuning curves (the four biological ones and label specificity) and for each model instantiation, distributions of test scores were computed (Figure 5, 7).

When plotting comparisons between different types of models (ART, TDT, and controls), the confidence interval for the mean (CLM) using an *α* = 5% significance level based on the t-statistic was displayed.

#### Label tuning (selectivity index)

The networks’ ability to solve the proprioceptive task poses the question if individual neurons serve as character detectors. To this end, SVMs were fit with linear kernels using a one vs. rest strategy for multi-class classification based on the firing rate of each node, resulting in linear decision boundaries for each letter. Each individual SVM serves as a binary classifier for the trajectory belonging to a certain character or not, based on that neuron’s firing rates. For each SVM, auROC was calculated, giving a measure of how well the label can be determined based on the firing rate of an individual node alone. The label specificity of that node was then determined by taking the maximum over all characters. Finally, the auROC score was normalized into a selectivity index: 2 ((auROC) − 0.5)).

#### Position, direction, velocity, & acceleration

For the kinematic tuning curves the coefficient of determination *R*^2^ on the test set was used as the primary metric of evaluation. These tuning curves were fitted using ordinary least squares linear regression, with regularization proving unnecessary due to the high number of data points and the low number of parameters (2-3) in the models.

#### Position tuning

Position 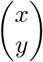 is initially defined with respect to the center of the workspace. For trajectories in a *horizontal* plane (workspace), a position vector was defined with respect to the starting position 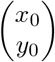 of each trace, 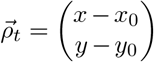. This was also represented in polar coordinates 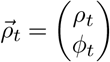, where *φ*_*t*_ ∈ (− *π, π*] is the angle measured with the counterclockwise direction defined as positive between the position vector and the vector 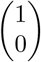, i.e. the vector extending away from the body, and 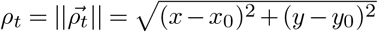. Positional tuning of the neural activity *N* of node *ν* was evaluated by fitting models both using Cartesian coordinates,

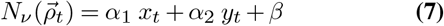

as well as polar ones,

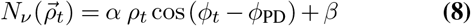

where *φ*_PD_ is a parameter representing a neuron’s preferred direction for position. For trajectories in the *vertical* plane, all definitions are equivalent, but with coordinates (*y, z*)^*T*^.

#### Direction

In order to examine the strength of kinematic tuning, tuning curves relating direction, velocity, and acceleration to neural activity were fitted. Since all trajectories take place either in a horizontal or vertical plane, the instantaneous velocity vector at time *t* can be described in two components as 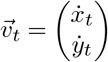 or (*y, z*)^*T*^ for trajectories in a vertical plane, or alternately in polar coordinates, 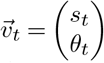, with *θ*_*t*_ ∈ (−*π, π*] representing the angle between the velocity vector and the x-axis, and 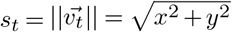 representing the speed.

First, a tuning curve was fit that excludes the magnitude of velocity but focuses on the instantaneous direction, putting the angle of the polar representation of velocity *θ*_*t*_ in relation to each neuron’s preferred direction *θ*_PD_.

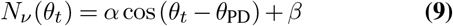

To fit this model, 9 was re-expressed as a simple linear sum using the cosine sum and difference formula cos(*α* + *β*) = cos *α* cos *β* − sin *α* sin *β*, a reformulation that eases the computational burden of the analysis significantly (70). In this formulation, the equation for directional tuning becomes:

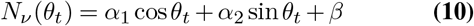

The preferred direction *θ*_PD_ is now contained in the in the coefficients *α*_1_ = *α* cos *θ*_PD_ and *α*_2_ = *α* sin *θ*_PD_.

The quality of fit of this type of tuning curve was visualized using polar scatter plots in which the angle of the data point corresponds to the angle *θ* in the polar representation of velocity and the radius corresponds to the node’s activation. In the figures the direction of movement was defined so that 0^*°*^ (Y) corresponds to movement to the right of the body and progressing counterclockwise, a movement straight (“forward”) away from the body corresponds to 90^*°*^ (X) (Figures 5A, B; 7A, B, C).

#### Speed

Two linear models for activity *N* at a node *ν* for velocity were fit.

The first is based on its magnitude, speed,

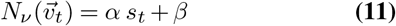

#### Velocity

The second velocity-based tuning curve factors in both directional and speed components:

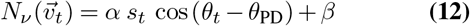

The quality of fit of this type of tuning curve was visualized using polar filled contour plots in which the angle of the data point corresponds to the angle *θ* in the polar representation of velocity, the radius corresponds to the speed, and the node’s activation is represented by the height. For the visualizations (Figure 5B), to cover the whole range of angle and radius given a finite number of samples, the activation was first linearly interpolated. Then, missing regions were filled in using nearest neighbor interpolation. Finally, the contour was smoothed using a Gaussian filter.

#### Acceleration

Acceleration is defined analogously to velocity by 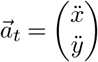 and 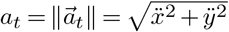. A simple linear relationship with acceleration magnitude was tested:

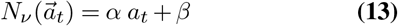

In subsequent analyses, scores were excluded if they were equal to 1 (indicating a dead neuron whose output was constant) or if they were less than -0.1 (indicating a fit that did not converge).

#### Classification of neurons into different types

The neurons were classified as belonging to a certain type if the corresponding kinematic model yielded a test-*R*^2^ *>* 0.2. Seven different model types were evaluated:

1. Direction tuning
2. Velocity tuning
3. Direction & velocity tuning
4. Position (Cartesian)
5. Position (Polar)
6. Acceleration tuning
7. Label specificity

These were treated as distinct classes for the purposes of classification (Figure 5C).

### Population Decoding Analysis

We also performed population-level decoding analysis for the kinematic tuning curve types (Figure S4, S8, S7). The same data sets were used as for the single cell encoding analysis, except with switched predictors and targets. The firing rates of all neurons in a hidden layer at a single time point were jointly used as predictors for the kinematic variable at the center of the receptive field at the corresponding input layer.

This analysis was repeated for each of the following kinematic variables:

1. Direction
2. Speed
3. X Position (Cartesian)
4. Y Position (Cartesian)

For each of these, the accuracy was evaluate using r2 score. The encoding strength for X and Y position in Cartesian coordinates was additionally jointly evaluated by calculating the average distance between true and predicted points of the trajectory. To prevent overfitting, ridge regularization was used with a regularization strength of *α* = 1.

### Distribution of Preferred Directions

Higher order features of the models were also evaluated and compared between the trained models and their controls. The first property was the distribution of preferred directions fit for all horizontal planes in each layer. If a neuron’s direction-only tuning yields a test-*R*^2^ *>* 0.2, its preferred direction was included in the distribution. Within a layer, the preferred direction of all neurons was binned into 18 equidistant intervals (Figure 7B,C) in order to enable a direct comparison with the findings by Prud’homme and Kalaska (14). They found that the preferred direction of tuning curves was relatively evenly spread in S1 (Figure 7A); our analysis showed that this was not the case for muscle spindles (Figure 7B). Thus, we formed the hypothesis that the preferred directions in the trained networks was more uniform in the trained networks than in the random ones. For quantification, absolute deviation from uniformity was used as a metric. To calculate this metric, the deviation from the mean height of a bin in the circular histograms was calculated for each angular bin. Then, the absolute value of this deviation was summed over all bins. We then normalize the result by the number of significantly directionally tuned neurons in a layer, and compare the result for the trained and control networks (Figure 7D).

### Preferred direction invariance

We also hypothesized that the representation in the trained network would be more invariant across different horizontal and vertical planes, respectively. To test this, directional tuning curves were fit for each individual plane. A central plane was chosen as a basis of comparison (plane at *z* = 0 for the horizontal planes and at *x* = 30 for vertical). Changes in preferred direction of neurons are shown for spindles (Figure S9A), as well as for neurons of layer 5 of one instantiation of the trained and control spatial-temporal model (Figure S9B). Generalization was then evaluated as follows: for neurons with *R*^2^ *>* 0.2, the average deviation of the neurons’ preferred directions over all different planes from those in the central plane was summed up and normalized by the number of planes and neurons, yielding a total measure for the neurons’ consistency in preferred direction in any given layer (vertical: Figure 7F; horizontal: Figure 7E). If a plane had fewer than three directionally tuned neurons, its results were excluded.

### Statistical Testing

To test whether differences were statistically significant between trained and control models paired t-tests were used with a pre-set significance level of *α* = 0.05.

## Supporting information

Supplementary Video Video depicts the OpenSim model being passively moved to match the human-drawn character "a" for three variants.

## Software

We used the scientific Python stack (python.org): Numpy, Pandas, Matplotlib, SciPy (71) and scikit-learn (66). Open-Sim (34, 62, 63) was used for biomechanics simulations and Tensorflow was used for constructing and training the neural network models (72).

## Code and data

Code and data will be shared upon publication.

## Acknowledgments

We are grateful to Alessandro Marin Vargas, Alberto Chiappa, Andy Bennetto and Bartek Borzyszkowski for comments on an earlier version of the manuscript. K.J.S. Current Address: Department of Experimental Psychology, University of Oxford, United Kingdom.

---

**Figure S1.**
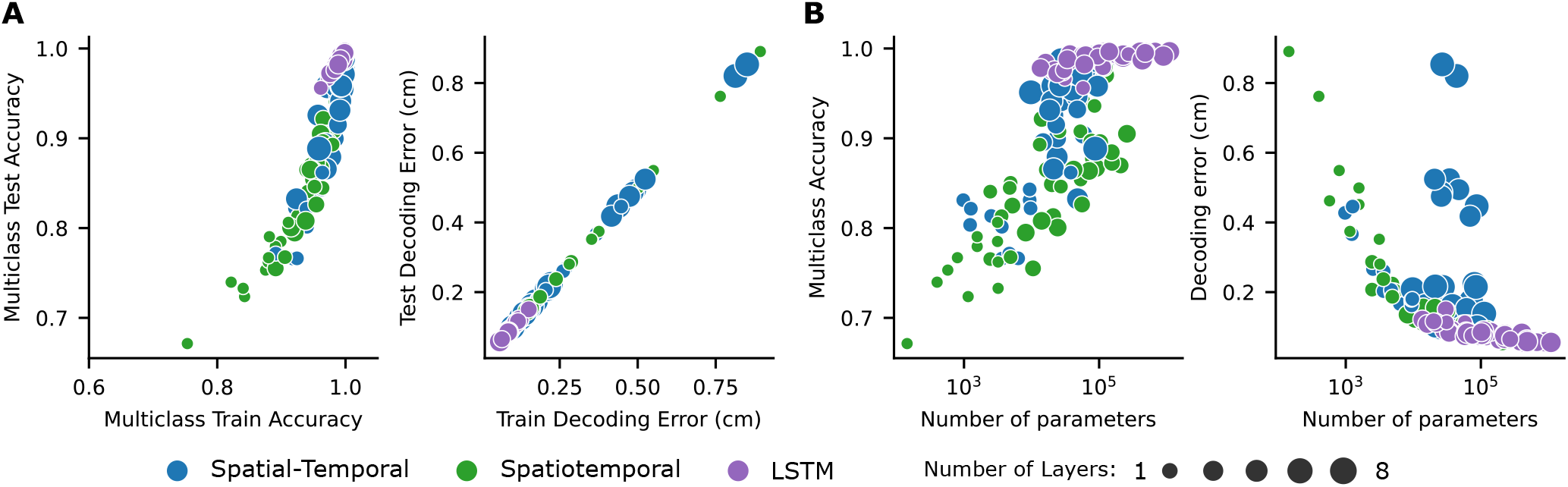
Network Performance. (**A**) Training vs. test performance for all networks. Shallower networks tend to overfit more. (**B**) Performance of networks is plotted against the all parameters of the networks. Note: parameters of the final (fully connected) layer are not counted.

**Figure S2.**
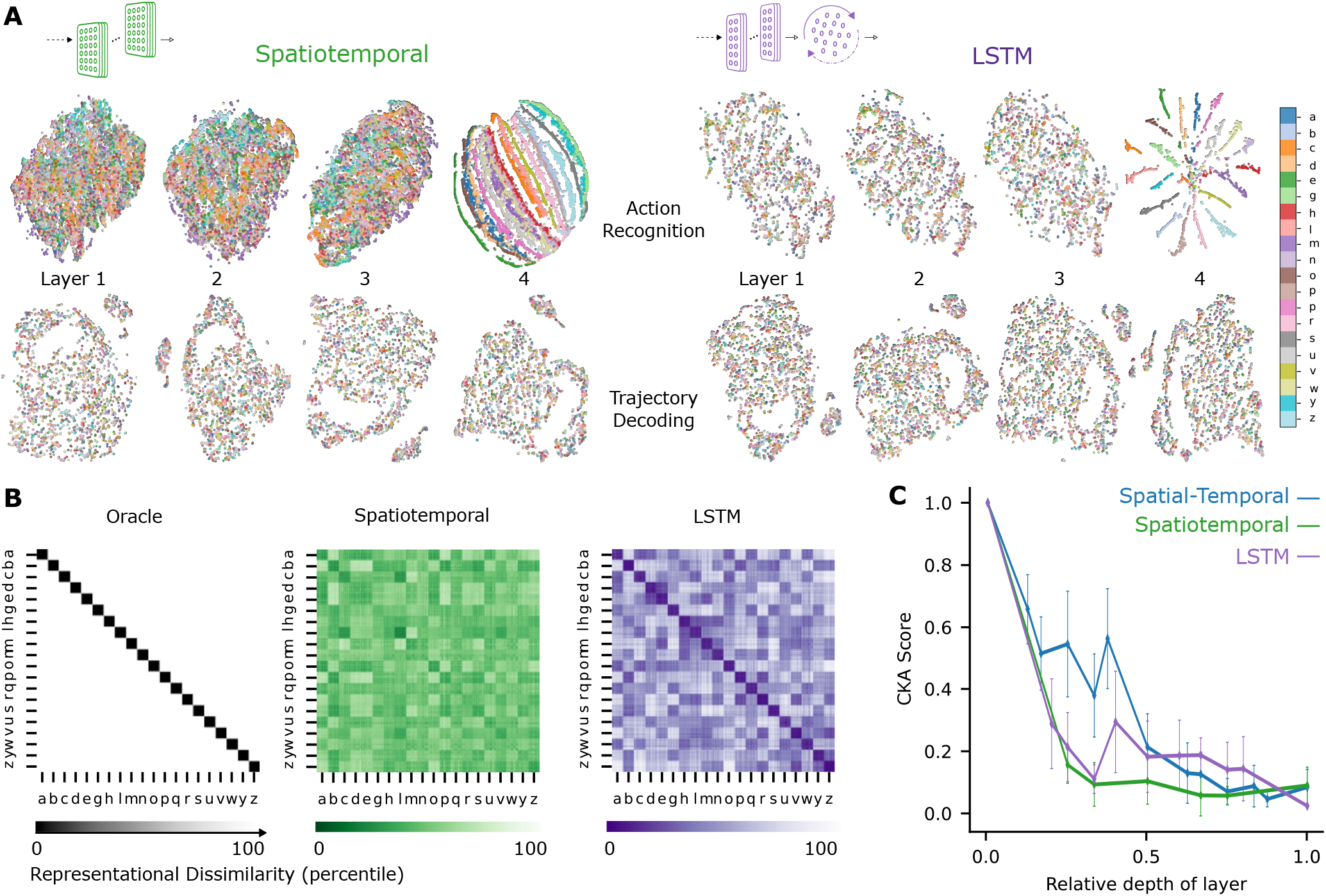
Extended analysis of network models. (**A**) t-distributed stochastic neighbor embedding (t-SNE) embedding for each layer of the best spatiotemporal and LSTM model. Each data point is a random stimulus sample (N=4,000, 200 per character). (**B**) Representational Dissimilarity Matrices (RDM) of an ideal observer “Oracle”, which by definition has low dissimilarity for different samples of the same character and high dissimilarity for different samples of different characters. Character level representation are calculated through percentile representational dissimilarity matrices for proprioceptive inputs and final layer features of one instantiation of the best performing spatiotemporal and LSTM model trained on recognition task.(**C**) CKA between models trained on recognition vs decoding for all network types (*N* = 50 per network type).

**Figure S3.**
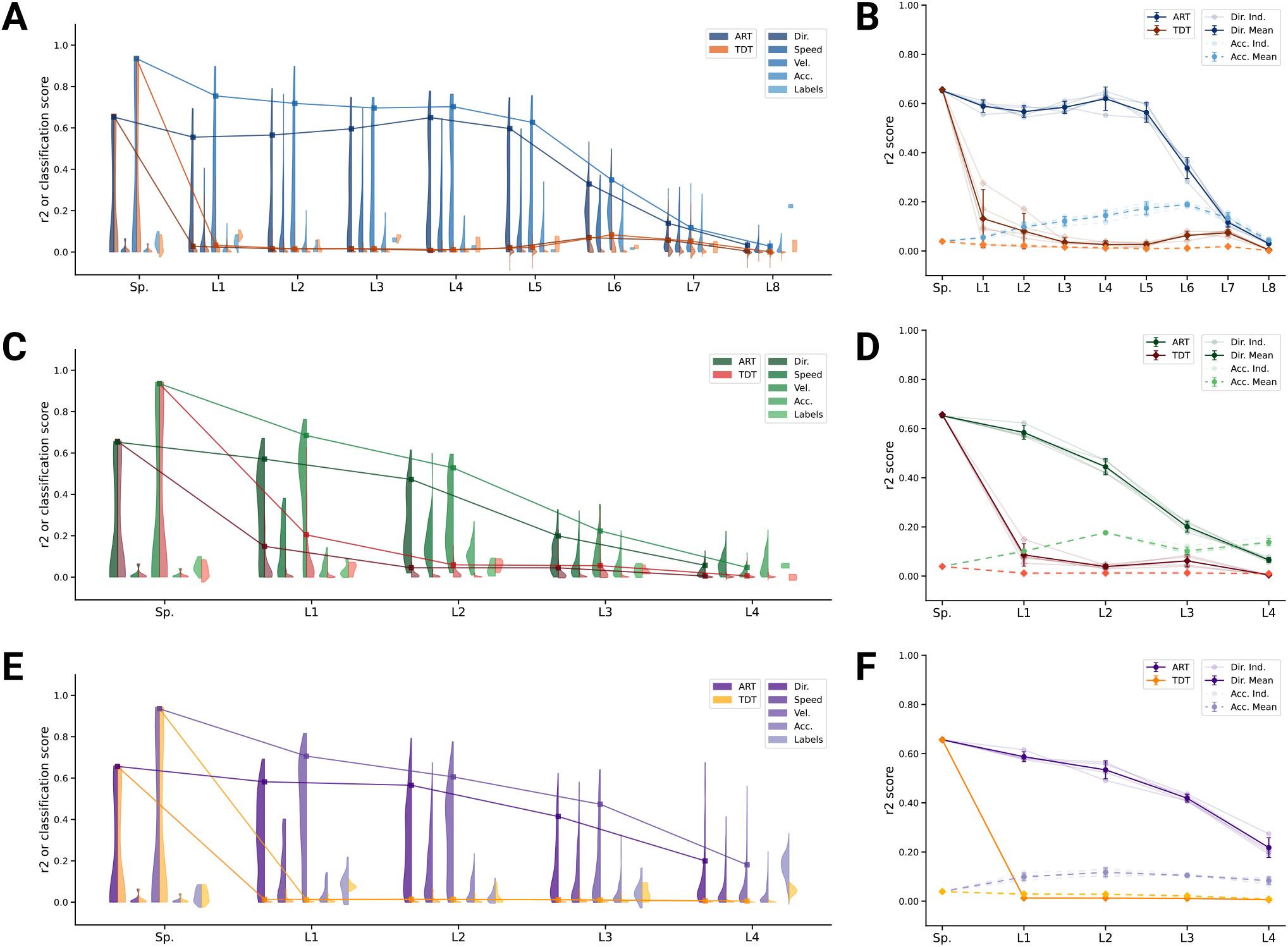
Kinematic tuning of single neurons. (**A**) For an example spatial-temporal model instantiation, the distribution of test *R*^2^ scores for both the ART- and TDT-trained models are shown, for direction, speed, velocity, acceleration, and labels. (8/3890 scores excluded over all layers for ART-trained, 284/3890 for TDT-trained; see Methods). (**B**) The individual traces (faint) as well as the means (dark) of 90%-quantiles over all five model instantiations of models trained on action recognition and trajectory decoding are shown for direction tuning (solid line) and acceleration tuning (dashed line). (**C**, D) Same as A,B but for the spatiotemporal model. (0/2330 scores excluded for ART-trained, 132/2330 for TDT; see Methods). (**E**, F) Same as A,B but for the LSTM model.(4/6530 scores excluded for ART-trained, 1024/6530 for TDT-trained; see Methods).

**Figure S4.**
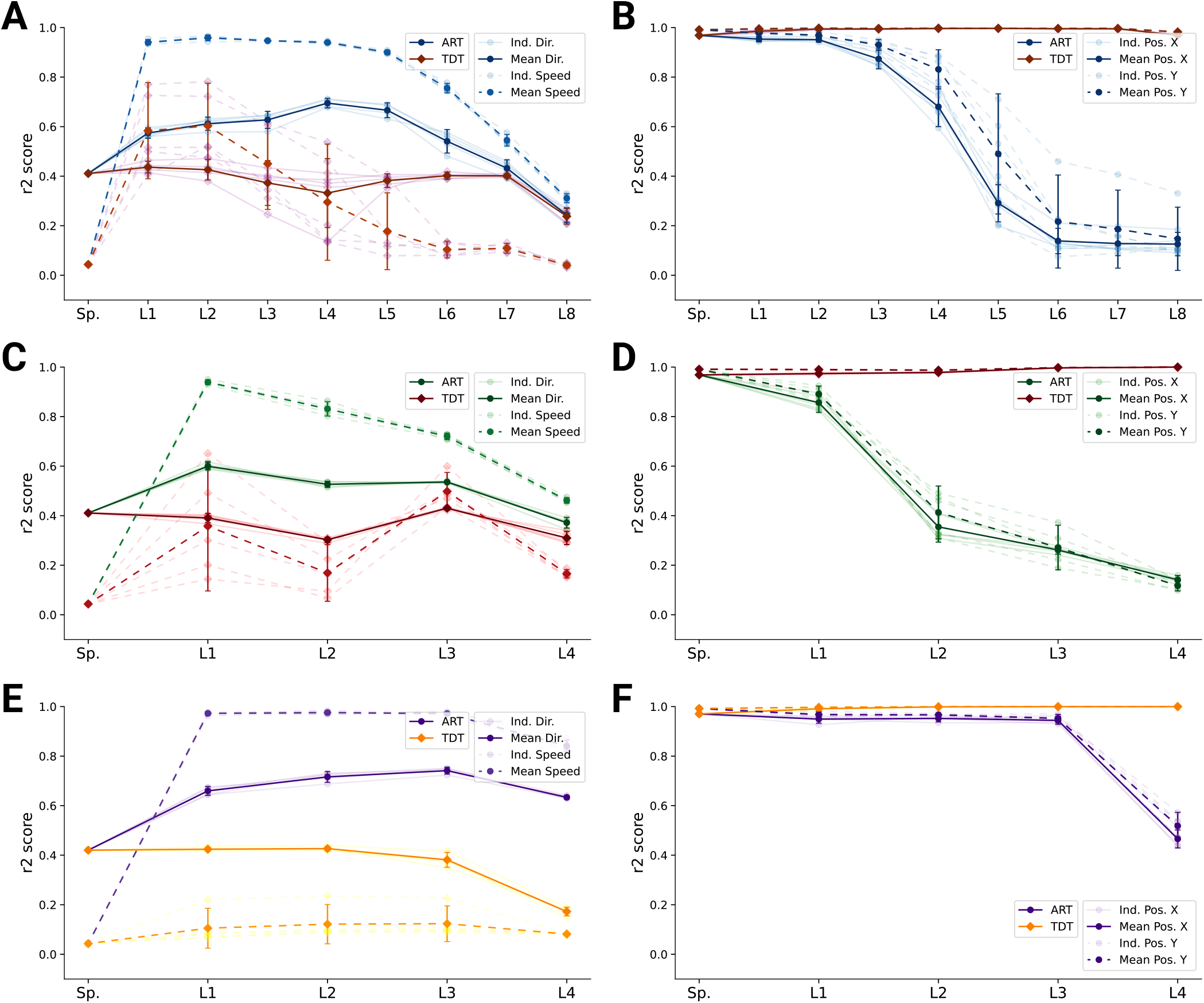
Population decoding analysis of ART vs. TDT models. (**A**) Population decoding of speed (*light*) and direction (*dark*) for spatial-temporal models for the ART- and TDT-trained models. The faint line shows the *R*^2^ score for an individual model; the dark one the mean over all instantiations (*N* = 5). (**B**) Population decoding of endeffector position (X and Y coordinates) for spatial temporal models. The faint line shows the *R*^2^ score for an individual model; the dark one the mean over all instantiations (*N* = 5). (**C**) Same as A but for spatiotemporal models. (**D**) Same as B but for spatiotemporal models. (**E**) Same as A but for LSTM models. (**F**) Same as B but for LSTM models.

**Figure S5.**
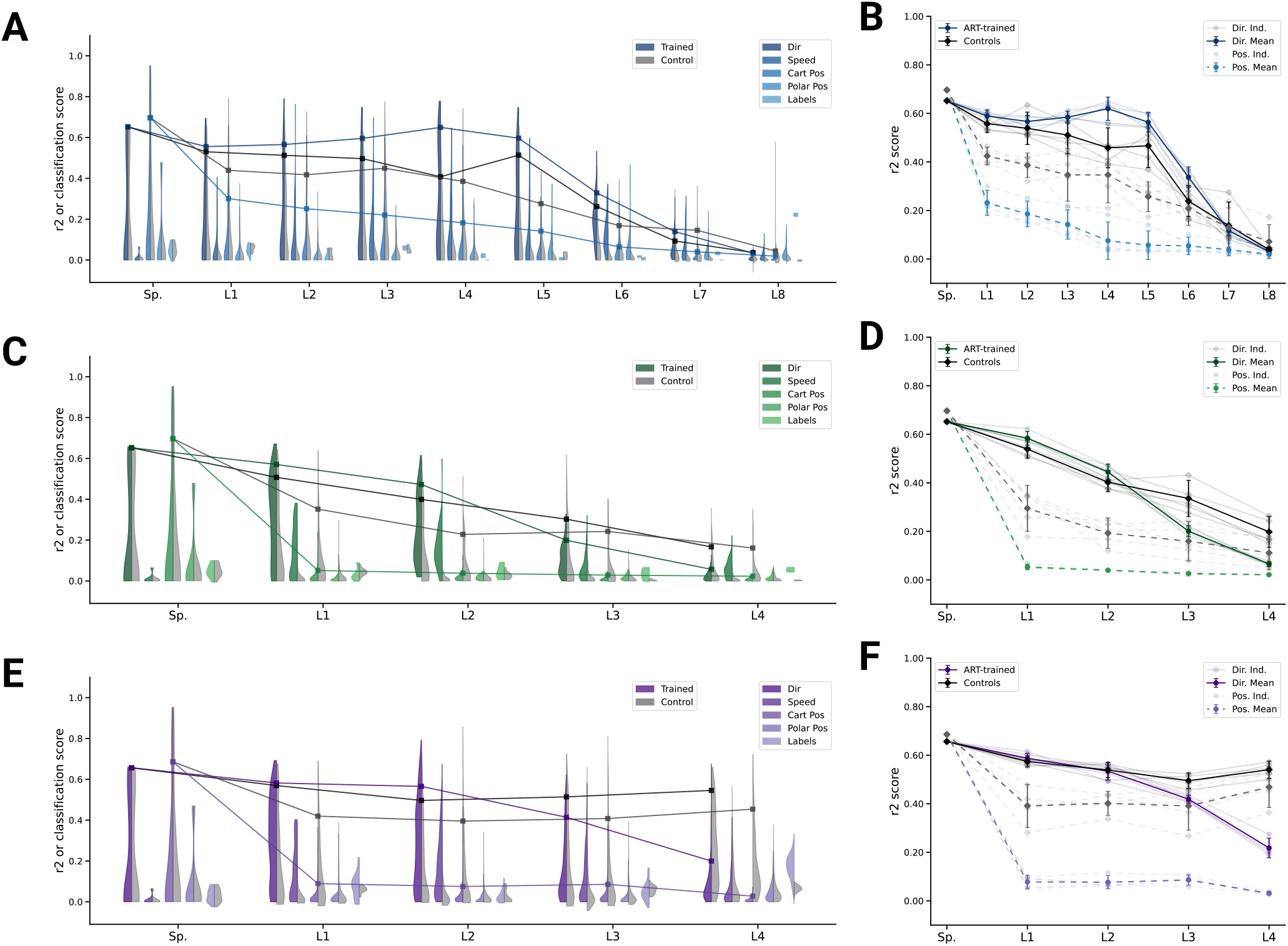
Analysis of single unit tuning properties for ART-trained models and controls. (**A**) For an example instantiation of the top-performing spatial-temporal model, the distribution of test *R*^2^ scores for both the trained and control model are shown, for five kinds of kinematic tuning for each layer: direction tuning, speed tuning, Cartesian position tuning, polar position tuning, and label-specificity. The solid line connects the 90%-quantiles of two of the tuning curve types, direction tuning (dark) and position tuning (light). (8/3890 scores excluded summed over all layers for ART-trained, 294/3890 for controls; see Methods). (**B**) The means of 90%-quantiles over all five model instantiations of models trained on action recognition and trajectory decoding are shown for direction tuning (dark) and position tuning (light). 95%-confidence intervals are shown over instantiations (*N* = 5). The same plot as in (A) but for the top-performing spatiotemporal model (133/2330 scores excluded for ART-trained, 60/2330 for the control; see Methods). (**D**) The same plot as B, for the spatiotemporal model. (**E**) The same plot as in (A) but for the top-performing LSTM model (4/6530 scores excluded for ART-trained, 328/6530 for the control; see Methods). (**F**) The same plot as B, for the LSTM model.

**Figure S6.**
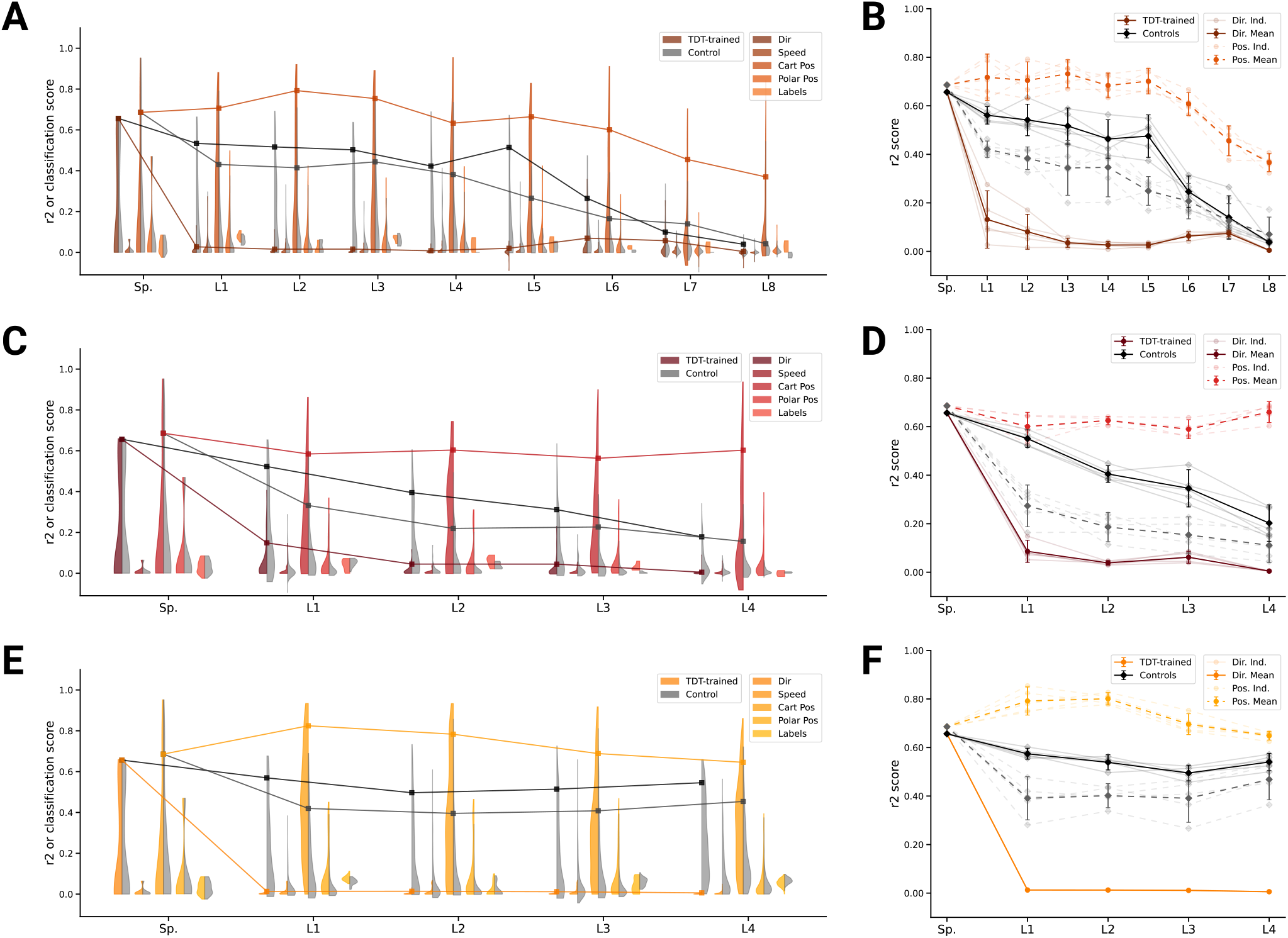
Analysis of single unit tuning properties for TDT-trained models and controls. (**A**) For an example instantiation of the top-performing spatial-temporal model, the distribution of test *R*^2^ scores for both the trained and control model are shown, for five kinds of kinematic tuning for each layer: direction tuning, speed tuning, Cartesian position tuning, polar position tuning, and label-specificity. The solid line connects the 90%-quantiles of two of the tuning curve types, direction tuning (dark) and position tuning (light). (287/3890 scores excluded summed over all layers for TDT-trained, 369/3890 for controls; see Methods). (**B**) The means of 90%-quantiles over all five model instantiations of models trained on action recognition and trajectory decoding are shown for direction tuning (dark) and position tuning (light). 95%-confidence intervals are shown over instantiations (*N* = 5). (**C**) The same plot as in (A) but for the top-performing spatiotemporal model (133/2330 scores excluded for TDT-trained, 60/2330 for the control; see Methods). (**D**) The same plot as B, for the spatiotemporal model. (**E**) The same plot as in (A) but for the top-performing LSTM model (1024/6530 scores excluded for TDT-trained, 328/6530 for the control; see Methods). (**F**) The same plot as B, for the LSTM model.

**Figure S7.**
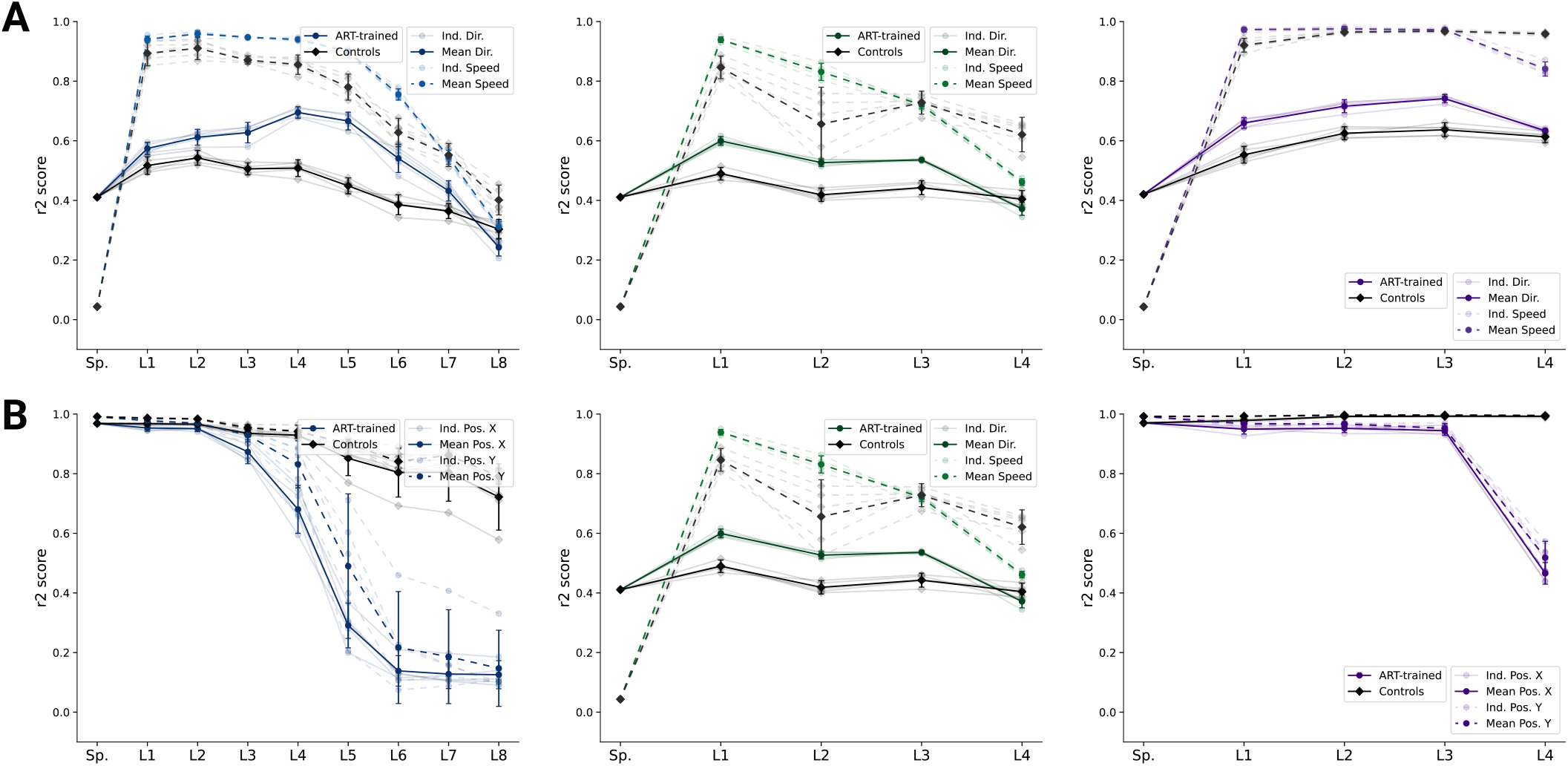
Analysis of population decoding for ART-trained and control models. (**A**) Population decoding of speed (*light*) and direction (*dark*) for the ART-trained and control for spatial-temporal models (*left*), spatiotemporal (*middle*) and LSTM (*right*) models. The faint line shows the *R*^2^ score for an individual model; the dark one the mean over all instantiations (*N* = 5). (**B**) Population decoding of endeffector position (X and Y coordinates) for spatial temporal models. The faint line shows the *R*^2^ score for an individual model; the dark one the mean over all instantiations (*N* = 5).

**Figure S8.**
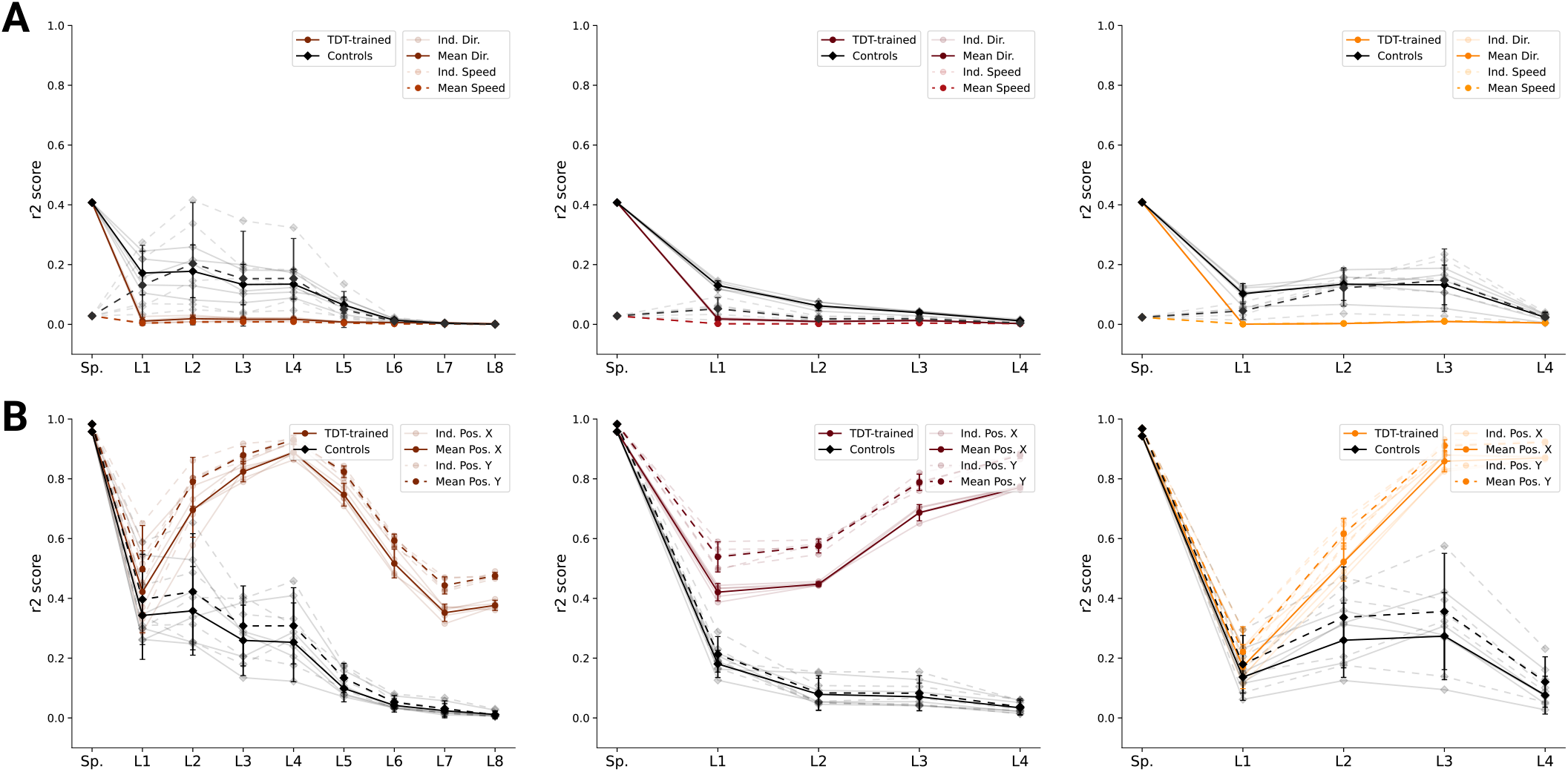
Analysis of population decoding for TDT-trained and control models. (**A**) Population decoding of speed (*light*) and direction (*dark*) for the TDT-trained and control for spatial-temporal models (*left*), spatiotemporal (*middle*) and LSTM (*right*) models. The faint line shows the *R*^2^ score for an individual model; the dark one the mean over all instantiations (*N* = 5). (**B**) Population decoding of endeffector position (X and Y coordinates) for spatial-temporal models (*left*), spatiotemporal (*middle*) and LSTM (*right*) models. The faint line shows the *R*^2^ score for an individual model; the dark one the mean over all instantiations (*N* = 5).

**Figure S9.**
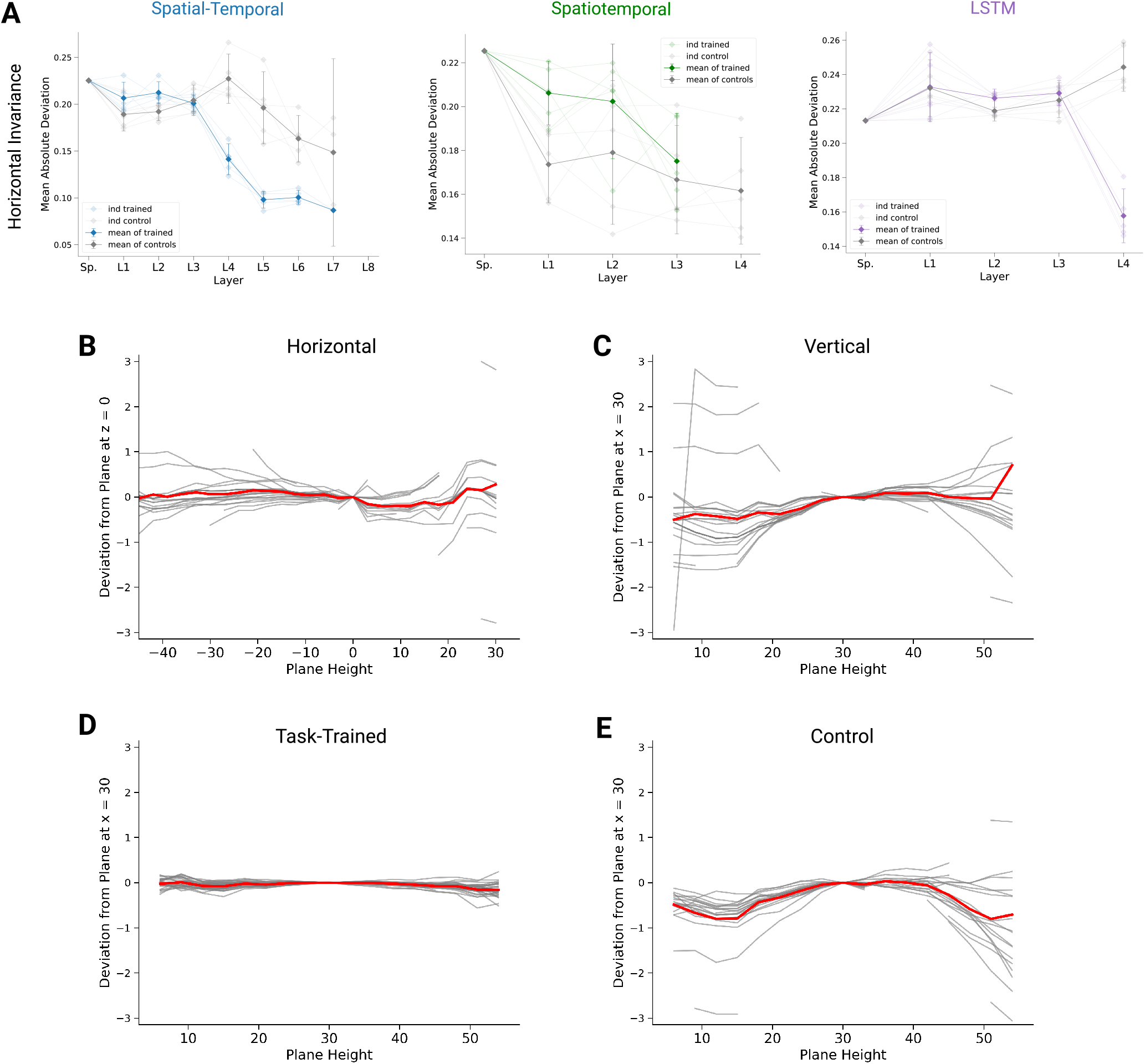
Invariance of preferred orientations. (**A**) For quantifying invariance we calculated mean absolute deviation in preferred orientation for units from a central plane at *z* = 0 to each other horizontal plane (for units with *R*^2^ *>* 0.2). Results are shown for each instantiation (*N* = 5, faint lines) for trained and control models plus mean (solid) and 95%-confidence intervals over instantiations (*N* = 5) for the spatial-temporal *(left)*, spatiotemporal *(right)*, and LSTM *(right)* networks. Note that there is no data for layer 4 of the trained spatiotemporal model, as it has no direction selective units (*R*^2^ *>* 0.2). (**B**) Deviation in preferred direction for individual spindles (N=25). The preferred directions are fit for each plane and displayed in relation to a central horizontal (*left*) and vertical plane (*right*). Individual gray lines are for all units (spindles) with *R*^2^ *>* 0.2, the thick red line marks the mean. (**C**) Same as B, but for direction tuning in vertical planes for units in layer 5 of one instantiation of the best spatial-temporal model for the trained (*left*) and control model (*right*). Individual gray lines are for units with *R*^2^ *>* 0.2, and the red line is the plane-wise mean. (**D**) Same as in B but for layer 5 of the trained spatial-temporal network. (**E**) Same as in D but for layer 5 of the corresponding control network.

**Figure S10.**
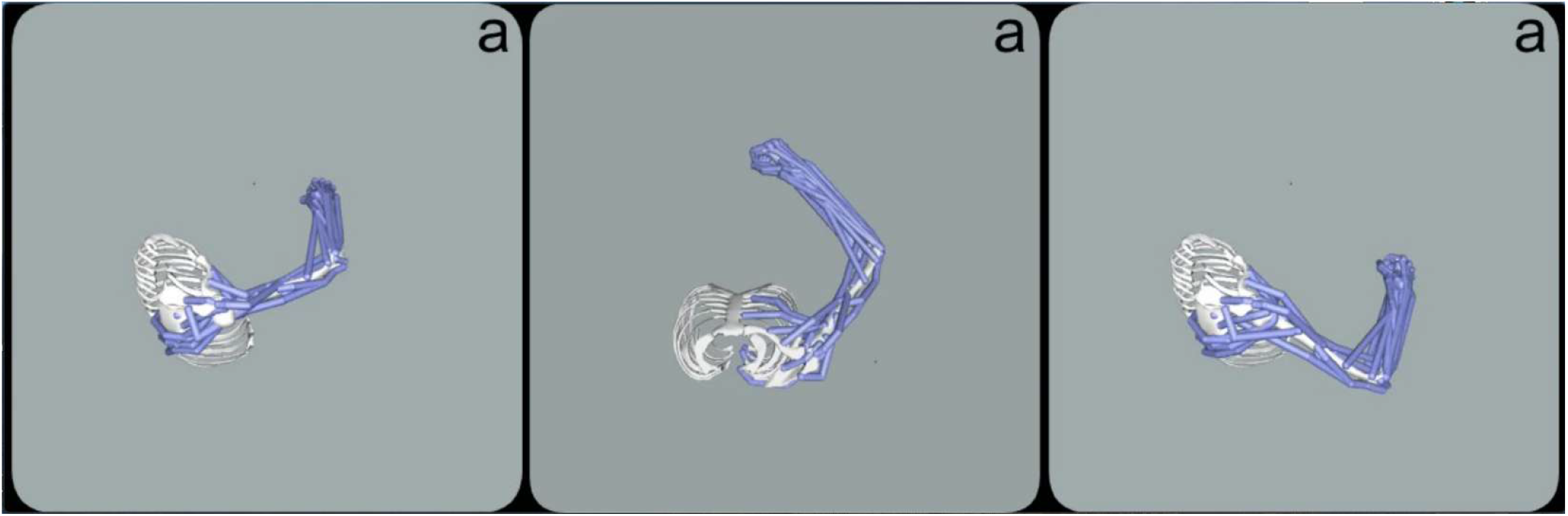
Supplementary Video. Video depicts the OpenSim model being passively moved to match the human-drawn character “a” for three different variants; drawn vertically (left, right) and horizontally (middle).

## References

1. R Chris Miall, Nick M Kitchen, Se-Ho Nam, Hannah Lefumat, Alix G Renault, Kristin Ørstavik, Jonathan D Cole, and Fabrice R Sarlegna. Proprioceptive loss and the perception, control and learning of arm movements in humans: evidence from sensory neuronopathy. Experimental brain research, pages 1–19, 2018.

2. Uwe Proske and Simon C Gandevia. The Proprioceptive Senses: Their Roles in Signaling Body Shape, Body Position and Movement, and Muscle Force. Physiological Reviews, 92(4):1651–1697, 2012. ISSN 0031-9333. doi: 10.1152/physrev.00048.2011.

3. Benoit P Delhaye, Katie H Long, and Sliman J Bensmaia. Neural basis of touch and proprioception in primate cortex. Comprehensive Physiology, 8(4):1575, 2018.

4. Francis J Clark, Curt R Burgess, James W Chapin, and WT Lipscomb. Role of intramuscular receptors in the awareness of limb position. Journal of Neurophysiology, 54(6):1529–1540, 1985.

5. Peter BC Matthews. The response of de-efferented muscle spindle receptors to stretching at different velocities. The Journal of Physiology, 168(3):660–678, 1963.

6. Peter BC Matthews. Muscle spindles: their messages and their fusimotor supply. Handbook of physiology: I. The nervous system. American Physiological Society.[AGF], 1981.

7. Nikolai A Bernstein. The co-ordination and regulation of movements, volume 1. Oxford, New York, Pergamon Press, 1967.

8. Gianfranco Bosco, A. Rankin, and Richard E Poppele. Representation of passive hindlimb postures in cat spinocerebellar activity. J Neurophysiol, 76(2):715–26, 1996. ISSN 0022-3077.

9. John C. Tuthill and Eiman Azim. Proprioception. Current Biology, 28(5):R193–R194, 2018. ISSN 09609822. doi: 10.1016/j.cub.2018.01.064.

10. Joseph T Francis, Shaohua Xu, and John K Chapin. Proprioceptive and cutaneous representations in the rat ventral posterolateral thalamus. Journal of neurophysiology, 99(5):2291–2304, 2008.

11. James M Goodman, Gregg A Tabot, Alex S Lee, Aneesha K Suresh, Alexander T Rajan, Nicholas G Hatsopoulos, and Sliman Bensmaia. Postural representations of the hand in the primate sensorimotor cortex. Neuron, 104(5):1000–1009, 2019.

12. Raeed H Chowdhury, Joshua I Glaser, and Lee E Miller. Area 2 of primary somatosensory cortex encodes kinematics of the whole arm. Elife, 9, 2020.

13. Christoph Fromm and Edward V Evarts. Pyramidal tract neurons in somatosensory cortex: central and peripheral inputs during voluntary movement. Brain research, 238(1):186–191, 1982.

14. Michel JL Prud’Homme and John F Kalaska. Proprioceptive activity in primate primary somatosensory cortex during active arm reaching movements. Journal of neurophysiology, 72(5):2280–2301, 1994.

15. Mackenzie W Mathis, Alexander Mathis, and Naoshige Uchida. Somatosensory cortex plays an essential role in forelimb motor adaptation in mice. Neuron, 93:p1493–1503.e6, 2017.

16. Neeraj Kumar, Timothy F Manning, and David J Ostry. Somatosensory cortex participates in the consolidation of human motor memory. PLoS biology, 17(10), 2019.

17. Uwe Proske and Simon C Gandevia. The proprioceptive senses: their roles in signaling body shape, body position and movement, and muscle force. Physiological reviews, 92(4):1651–1697, 2012.

18. Michael SA Graziano. Ethological action maps: a paradigm shift for the motor cortex. Trends in cognitive sciences, 20(2):121–132, 2016.

19. Olga Russakovsky, Jia Deng, Hao Su, Jonathan Krause, Sanjeev Satheesh, Sean Ma, Zhiheng Huang, Andrej Karpathy, Aditya Khosla, Michael Bernstein, et al. Imagenet large scale visual recognition challenge. International Journal of Computer Vision, 115(3): 211–252, 2015.

20. Seyed-Mahdi Khaligh-Razavi and Nikolaus Kriegeskorte. Deep supervised, but not unsupervised, models may explain it cortical representation. PLoS computational biology, 10(11):e1003915, 2014.

21. Daniel LK Yamins, Ha Hong, Charles F Cadieu, Ethan A Solomon, Darren Seibert, and James J DiCarlo. Performance-optimized hierarchical models predict neural responses in higher visual cortex. Proceedings of the National Academy of Sciences, 111(23):8619–8624, 2014.

22. Radoslaw Martin Cichy, Aditya Khosla, Dimitrios Pantazis, Antonio Torralba, and Aude Oliva. Comparison of deep neural networks to spatio-temporal cortical dynamics of human visual object recognition reveals hierarchical correspondence. Scientific reports, 6: 27755, 2016.

23. Daniel LK Yamins and James J DiCarlo. Using goal-driven deep learning models to understand sensory cortex. Nature neuroscience, 19(3):356, 2016.

24. Martin Schrimpf, Jonas Kubilius, Ha Hong, Najib J Majaj, Rishi Rajalingham, Elias B Issa, Kohitij Kar, Pouya Bashivan, Jonathan Prescott-Roy, Kailyn Schmidt, et al. Brain-score: Which artificial neural network for object recognition is most brain-like? BioRxiv, page 407007, 2018.

25. Santiago A Cadena, George H Denfield, Edgar Y Walker, Leon A Gatys, Andreas S Tolias, Matthias Bethge, and Alexander S Ecker. Deep convolutional models improve predictions of macaque v1 responses to natural images. PLoS computational biology, 15(4): e1006897, 2019.

26. Katherine R Storrs, Tim C Kietzmann, Alexander Walther, Johannes Mehrer, and Nikolaus Kriegeskorte. Diverse deep neural networks all predict human inferior temporal cortex well, after training and fitting. Journal of Cognitive Neuroscience, 33(10):2044–2064, 2021.

27. Blake A Richards, Timothy P Lillicrap, Philippe Beaudoin, Yoshua Bengio, Rafal Bogacz, Amelia Christensen, Claudia Clopath, Rui Ponte Costa, Archy de Berker, Surya Ganguli, et al. A deep learning framework for neuroscience. Nature Neuroscience, 22(11): 1761–1770, 2019.

28. Andrew Saxe, Stephanie Nelli, and Christopher Summerfield. If deep learning is the answer, then what is the question? arXiv preprint 2004.07580, 2020.

29. Chengxu Zhuang, Jonas Kubilius, Mitra JZ Hartmann, and Daniel L Yamins. Toward goal-driven neural network models for the rodent whisker-trigeminal system. In Advances in Neural Information Processing Systems, pages 2555–2565, 2017.

30. Subramanian Sundaram, Petr Kellnhofer, Yunzhu Li, Jun-Yan Zhu, Antonio Torralba, and Wojciech Matusik. Learning the signatures of the human grasp using a scalable tactile glove. Nature, 569(7758): 698–702, 2019.

31. Martin Haesemeyer, Alexander F Schier, and Florian Engert. Convergent temperature representations in artificial and biological neural networks. Neuron, 103(6):1123–1134, 2019.

32. Alexander JE Kell, Daniel LK Yamins, Erica N Shook, Sam V Norman-Haignere, and Josh H McDermott. A task-optimized neural network replicates human auditory behavior, predicts brain responses, and reveals a cortical processing hierarchy. Neuron, 98 (3):630–644, 2018.

33. Ben H Williams, Marc Toussaint, and Amos J Storkey. Extracting motion primitives from natural handwriting data. In International Conference on Artificial Neural Networks, pages 634–643. Springer, 2006.

34. Katherine R Saul, Xiao Hu, Craig M Goehler, Meghan E Vidt, Melissa Daly, Anca Velisar, and Wendy M Murray. Benchmarking of dynamic simulation predictions in two software platforms using an upper limb musculoskeletal model. Computer methods in biomechanics and biomedical engineering, 18(13):1445–1458, 2015.

35. Michael Dimitriou and Benoni B Edin. Discharges in human muscle receptor afferents during block grasping. Journal of Neuroscience, 28(48):12632–12642, 2008.

36. Michael Dimitriou and Benoni B Edin. Discharges in human muscle spindle afferents during a key-pressing task. The Journal of physiology, 586(22):5455–5470, 2008.

37. Dua Dheeru and Efi Karra Taniskidou. UCI machine learning repository, 2017.

38. Laurens van der Maaten and Geoffrey Hinton. Visualizing data using t-sne. Journal of machine learning research, 9(Nov):2579–2605, 2008.

39. Chih-Wei Hsu and Chih-Jen Lin. A comparison of methods for multiclass support vector machines. IEEE transactions on Neural Networks, 13(2):415–425, 2002.

40. Sébastien B Hausmann, Alessandro Marin Vargas, Alexander Mathis, and Mackenzie W Mathis. Measuring and modeling the motor system with machine learning. Current Opinion in Neurobiology, 70:11–23, 2021.

41. David E Rumelhart, Geoffrey E Hinton, James L McClelland, et al. A general framework for parallel distributed processing. Parallel distributed processing: Explorations in the microstructure of cognition, 1(45-76):26, 1986.

42. Yann LeCun, Léon Bottou, Yoshua Bengio, Patrick Haffner, et al. Gradient-based learning applied to document recognition. Proceedings of the IEEE, 86(11):2278–2324, 1998.

43. Sepp Hochreiter and Jürgen Schmidhuber. Long short-term memory. Neural computation, 9(8):1735–1780, 1997.

44. Shaojie Bai, J Zico Kolter, and Vladlen Koltun. An empirical evaluation of generic convolutional and recurrent networks for sequence modeling. arXiv preprint 1803.01271, 2018.

45. Diederik P Kingma and Jimmy Ba. Adam: A method for stochastic optimization. arXiv preprint 1412.6980, 2014.

46. Edith Ribot-Ciscar, Mikael Bergenheim, Frédéric Albert, and Jean-Pierre Roll. Proprioceptive population coding of limb position in humans. Experimental brain research, 149(4):512–519, 2003.

47. P. Kibleur, S. R. Tata, N. Greiner, S. Conti, B. Barra, K. Zhuang, M. Kaeser, A. Ijspeert, and M. Capogrosso. Spatiotemporal maps of proprioceptive inputs to the cervical spinal cord during three-dimensional reaching and grasping. IEEE Transactions on Neural Systems and Rehabilitation Engineering, pages 1–1, 2020.

48. Bernd Illing, Wulfram Gerstner, and Johanni Brea. Biologically plausible deep learning—but how far can we go with shallow networks? Neural Networks, 118:90–101, 2019.

49. Frederic Albert, Edith Ribot-Ciscar, Michel Fiocchi, Mikael Bergenheim, and Jean-Pierre Roll. Proprioceptive feedback in humans expresses motor invariants during writing. Experimental brain research, 164(2):242–249, 2005.

50. Kai J Sandbrink, Pranav Mamidanna, Claudio Michaelis, Mackenzie Weygandt Mathis, Matthias Bethge, and Alexander Mathis. Task-driven hierarchical deep neural network models of the proprioceptive pathway. bioRxiv, 2020.

51. Arthur Prochazka and Monica Gorassini. Ensemble firing of muscle afferents recorded during normal locomotion in cats. The Journal of physiology, 507(1):293–304, 1998.

52. Christopher Versteeg, Joshua M Rosenow, Sliman J Bensmaia, and Lee E Miller. Encoding of limb state by single neurons in the cuneate nucleus of awake monkeys. Journal of Neurophysiology, 2021.

53. Thomas Serre. Deep learning: the good, the bad, and the ugly. Annual Review of Vision Science, 5:399–426, 2019.

54. Andrew Jackson, Jaideep Mavoori, and Eberhard E Fetz. Correlations between the same motor cortex cells and arm muscles during a trained task, free behavior, and natural sleep in the macaque monkey. Journal of neurophysiology, 97(1):360–374, 2007.

55. Simon C Gandevia, Kathyrn M Refshauge, and David F Collins. Proprioception: peripheral inputs and perceptual interactions. In Sensorimotor control of movement and posture, pages 61–68. Springer, 2002.

56. Milana P Mileusnic, Ian E Brown, Ning Lan, and Gerald E Loeb. Mathematical models of proprioceptors. i. control and transduction in the muscle spindle. Journal of neurophysiology, 96(4):1772–1788, 2006.

57. Jean-Marc Aimonetti, Valérie Hospod, Jean-Pierre Roll, and Edith Ribot-Ciscar. Cutaneous afferents provide a neuronal population vector that encodes the orientation of human ankle movements. The Journal of physiology, 580(2):649–658, 2007.

58. Kyle P Blum, Boris Lamotte D’Incamps, Daniel Zytnicki, and Lena H Ting. Force encoding in muscle spindles during stretch of passive muscle. PLoS computational biology, 13(9):e1005767, 2017.

59. Rochelle Ackerley, Marie Chancel, Jean-Marc Aimonetti, Edith Ribot-Ciscar, and Anne Kavounoudias. Seeing your foot move changes muscle proprioceptive feedback. Eneuro, 6(2), 2019.

60. Katherine RS Holzbaur, Wendy M Murray, and Scott L Delp. A model of the upper extremity for simulating musculoskeletal surgery and analyzing neuromuscular control. Annals of biomedical engineering, 33(6):829–840, 2005.

61. Aaron D’Souza, Sethu Vijayakumar, and Stefan Schaal. Learning inverse kinematics. In Proceedings 2001 IEEE/RSJ International Conference on Intelligent Robots and Systems. Expanding the Societal Role of Robotics in the the Next Millennium (Cat. No. 01CH37180), volume 1, pages 298–303. IEEE, 2001.

62. Scott L Delp, Frank C Anderson, Allison S Arnold, Peter Loan, Ayman Habib, Chand T John, Eran Guendelman, and Darryl G Thelen. Opensim: open-source software to create and analyze dynamic simulations of movement. IEEE transactions on biomedical engineering, 54(11):1940–1950, 2007.

63. Ajay Seth, Michael Sherman, Jeffrey A Reinbolt, and Scott L Delp. Opensim: a musculoskeletal modeling and simulation framework for in silico investigations and exchange. Procedia Iutam, 2:212–232, 2011.

64. Vaughan G Macefield and Thomas P Knellwolf. Functional properties of human muscle spindles. Journal of neurophysiology, 2018.

65. Arthur Prochazka and Monica Gorassini. Models of ensemble firing of muscle spindle afferents recorded during normal locomotion in cats. The Journal of physiology, 507(1):277–291, 1998.

66. Fabian Pedregosa, Gaël Varoquaux, Alexandre Gramfort, Vincent Michel, Bertrand Thirion, Olivier Grisel, Mathieu Blondel, Peter Prettenhofer, Ron Weiss, Vincent Dubourg, et al. Scikit-learn: Machine learning in python. the Journal of machine Learning research, 12: 2825–2830, 2011.

67. Jimmy Lei Ba, Jamie Ryan Kiros, and Geoffrey E Hinton. Layer normalization. arXiv preprint 1607.06450, 2016.

68. Simon Kornblith, Mohammad Norouzi, Honglak Lee, and Geoffrey Hinton. Similarity of neural network representations revisited. arXiv preprint 1905.00414, 2019.

69. Nikolaus Kriegeskorte, Marieke Mur, and Peter A Bandettini. Representational similarity analysis-connecting the branches of systems neuroscience. Frontiers in systems neuroscience, 2:4, 2008.

70. Apostolos P Georgopoulos, John F Kalaska, Roberto Caminiti, and Joe T Massey. On the relations between the direction of two-dimensional arm movements and cell discharge in primate motor cortex. The Journal of neuroscience : the official journal of the Society for Neuroscience, 2 11:1527–37, 1982.

71. Pauli Virtanen, Ralf Gommers, Travis E Oliphant, Matt Haberland, Tyler Reddy, David Cournapeau, Evgeni Burovski, Pearu Peterson, Warren Weckesser, Jonathan Bright, et al. Scipy 1.0: fundamental algorithms for scientific computing in python. Nature methods, 17 (3):261–272, 2020.

72. Martín Abadi, Paul Barham, Jianmin Chen, Zhifeng Chen, Andy Davis, Jeffrey Dean, Matthieu Devin, Sanjay Ghemawat, Geoffrey Irving, Michael Isard, et al. Tensorflow: A system for large-scale machine learning. In 12th {USENIX} Symposium on Operating Systems Design and Implementation ({OSDI} 16), pages 265–283, 2016.

